# Time-Domain Diffuse Optical Tomography for Precision Neuroscience

**DOI:** 10.1101/2024.04.30.591765

**Authors:** Yaroslav Chekin, Dakota Decker, Hamid Dehghani, Julien Dubois, Ryan M. Field, Viswanath Gopalakrishnan, Erin M. Koch, Gabriel Lerner, Zahra M. Aghajan, Naomi Miller, Isai Olvera, Milin J. Patel, Katherine L. Perdue, Joshua Schmidt, Victor Szczepanski

**Affiliations:** Kernel, Culver City, CA 90232 USA

## Abstract

Recent years have witnessed a rise in research utilizing neuroimaging for precision neuromedicine, but clinical translation has been hindered by scalability and cost. Time Domain functional Near Infrared Spectroscopy (TD-fNIRS), the gold standard of optical neuroimaging techniques, offers a unique opportunity in this domain since it provides superior depth sensitivity and enables resolution of absolute properties unlike its continuous wave counterparts. However, current TD systems have limited commercial availability, slow sampling rates, and sparse head coverage. Our team has overcome the technical challenges involved in developing a whole-head time-domain diffuse optical tomography (TD-DOT) system. Here, we present the system characterization results using standardized protocols and compare them to the state-of-the-art. Furthermore, we showcase the system performance in retrieving cortical activation maps during standard hemodynamic, sensory, and motor tasks. A combination of the system performance, signal quality, and ease-of-use can enable future studies aimed at investigating TD-DOT clinical applications.

## Introduction

Neurocognitive and psychiatric disorders are highly prevalent and treatment expenditures are projected to increase in the coming decades (*1, 2*). Many emerging treatments for these disorders leverage novel mechanisms of action from scientific research for improved clinical results. However, clinical decision making around diagnosis, treatment selection, and treatment monitoring for these complex disorders remains a challenge. Quantitative brain data may be able to help clinicians make informed decisions for personalized care. As an example, evidence is building that non-invasive neuroimaging methods like functional magnetic resonance imaging (fMRI) (*3*–*6*) or electroencephalography (EEG) (*7*), may be instrumental for tailored depression treatments. In their current implementation, non-invasive neuroimaging assays are typically expensive and inaccessible, and may be contraindicated for certain patients. Methods to quantify the brain in a simple, quick, informative way at a point-of-care clinic are urgently needed.

Wearable functional near-infrared spectroscopy (fNIRS) systems have the potential to fill this gap (*8*). In the past 15 years, fNIRS systems have become more comfortable and capable, with higher spatial resolution, multimodal measurement capabilities, and standardized data collection and analysis procedures (*9*). With these improvements, wearable fNIRS systems enable measurements in situations inaccessible or inconvenient for other forms of neuroimaging, including during free ambulation (*10*), social interactions (*11*), and at point-of-care clinics.

While these improvements have allowed for an increased range of measurement situations, the underlying technology of most wearable fNIRS systems has remained the same: Continuous Wave (CW) light. CW systems have the benefit of being relatively inexpensive and straightforward, although systems designed for High-Density Diffuse Optical Tomography (HD-DOT) have a large number of optical sources and detectors which can increase cost and system complexity, as well as reduce sampling frequency (*12*). Alternatively, Time-Domain (TD-)fNIRS can go beyond traditional HD-DOT systems and interrogate the underlying tissue in greater detail due to its higher depth sensitivity (*13*). TD-fNIRS systems use short pulses of light and detectors capable of measuring single photons to capture a distribution of times of flight (DTOF) of photons. This fine-grained measurement capability allows TD-fNIRS systems to measure absolute optical properties of the underlying tissue including the absorption (μ_a_) and reduced scattering (μ_s_′) coefficients, instead of only measuring relative changes in light intensity like CW systems. The increased information from TD-fNIRS systems also allows for advanced signal processing methods to more heavily weight photons that arrive late in the DTOF in order to emphasize data from deeper in the head (i.e., from the brain). Despite these benefits, TD-fNIRS systems have limited commercial availability, and the available systems have slow sampling frequencies or limited coverage over the head.

Our team has been working to overcome the technical challenges involved in developing a portable, whole-head coverage, high-density, fast, and scalable TD-fNIRS system. We previously published on the characterization and validation of our prototype system, Kernel Flow1 (*14*). While the prototype system was used in several scientific studies (*13, 15, 16*), it had some limitations: there were gaps in spatial coverage over the head; pre-recorded instrument response functions (IRFs) were not sufficiently stable in time to ensure accuracy of absolute metrics; detectors had limited sensitivity; and the system had high power requirements. As such, we had confined our prior analyses to those performed in channel space and did not fully explore Flow1’s whole-head DOT capabilities.

In this paper, we present the first whole-head coverage Time-Domain Diffuse Optical Tomography (TD-DOT) system, Flow2. We first show that Flow2 compares favorably with limited channel count research-grade devices on key figures of merit agreed upon by the field, while extending the field-of-view to cover the whole head. We then demonstrate how it can be used to measure absolute concentrations of oxy- and deoxy-hemoglobin in the brain (during a breath hold challenge), and how it can be used to reconstruct focal brain activity (during a sensory and a motor task). Improvements in reconstruction accuracy for TD vs. CW data have been theoretically discussed (*17, 18*), and empirically demonstrated with limited coverage systems (*19*). Here we empirically show the benefit of whole-head TD over CW systems. In light of these characterization results and the scalability-by-design of the Flow2 device, we argue that it is well positioned to enable the translation of research findings from both fNIRS and fMRI literatures to clinical settings.

## Results

### Kernel Flow2: a wearable system for TD-DOT

Kernel Flow2 is a wearable time-domain fNIRS system that has been designed to be low-cost and compact while achieving dense channel coverage over the entire head (Fig. 1A). It has a modular design (Fig. 1B-F), with 40 modules spanning the entire headset. The device uses visible and near-infrared light (690 nm and 905 nm) to measure changes in blood oxygenation. It features more than 3500 source-detector pairs (counting sources and detectors that are less than 60 mm apart; or 2500 if limiting source-detector distance to 50 mm); the combination of information from channels that sample from the same brain region allows for spatial reconstruction of oxygenation changes (diffuse optical tomography). Further details about the design and engineering choices can be found in the Materials and Methods section.

**Figure 1.**
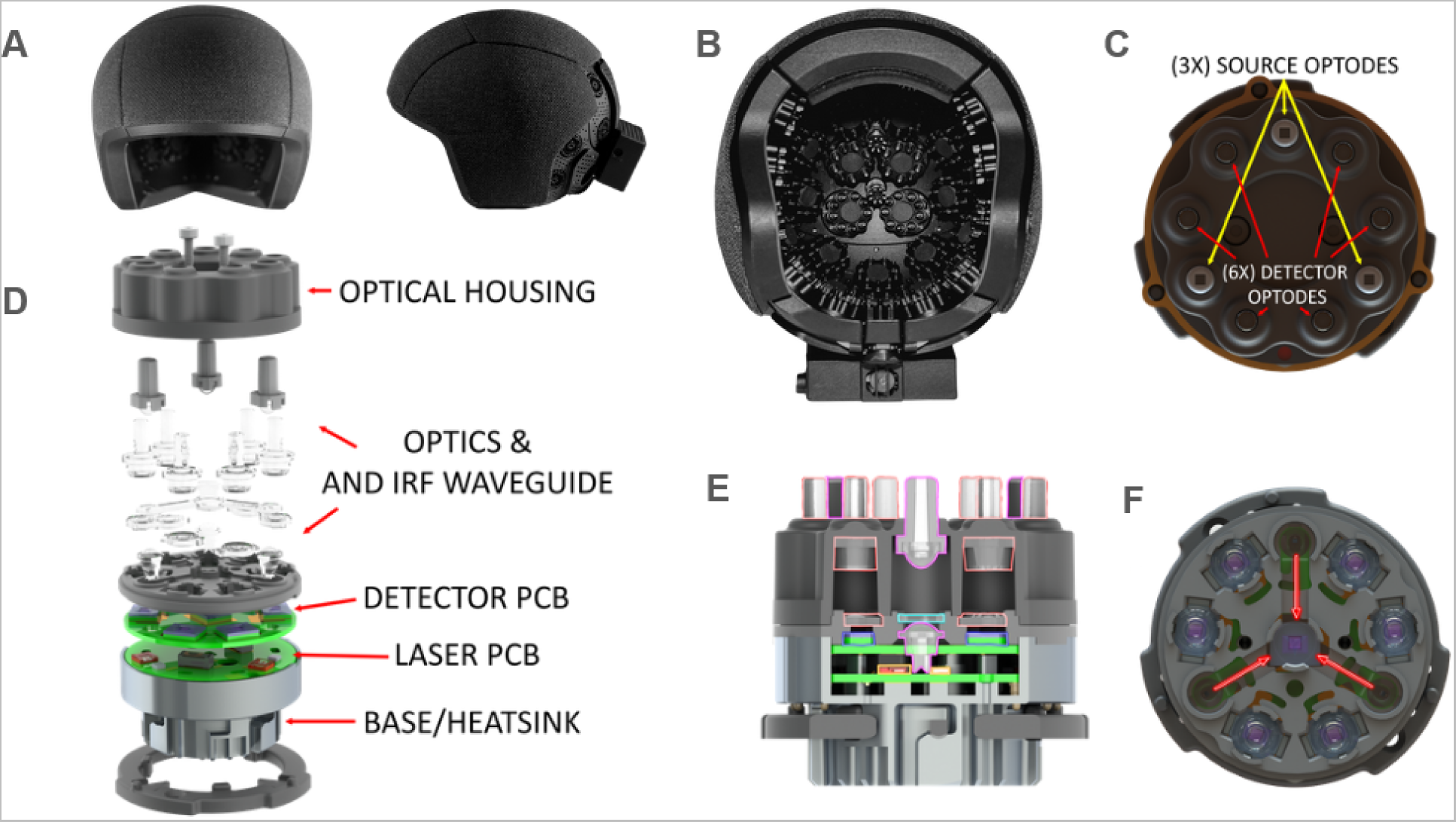
Kernel Flow2 headset provides modular coverage over the whole head with thousands of source-detector optical channels. **(A)** The Kernel Flow2 headset, exterior appearance. **(B)** The system consists of 40 modules: modules are organized into 7 headset rigid superstructure plates that cover the frontal, parietal, temporal, and occipital cortices. **(C)** Each Kernel Flow2 optical module consists of 3 dual-wavelength sources and 6 detectors located on a 13.5 mm radius from module center. An additional detector located in the center of the module continuously measures the IRF (see also E, F). The 3 source emission points are offset 120 degrees, with a detector located 37 degrees on either side of each source point. **(D)** Exploded view of the Flow2 module showing the details of all the module subassemblies, as labeled. **(E)** A cutout view of the optical module assembly, showing the source optics (magenta outline) consisting of a combined anti-prism and first element of the source optics and a combined exit lens and homogenizing light tunnel. The detector aspheric fresnel lenses are also visible (orange outline). **(F)** The IRF waveguide deflection design; this approach uses a specifically designed waveguide that sits in the optical path of the laser beams to transmit a fraction of the light from each source to the dedicated reference IRF detector.

### System performance assessment: BIP, MEDPHOT and nEUROPt protocols

The performance of a novel system must first be evaluated, with respect to systems of its class, using established protocols and well-defined figures of merit (*20*). Three characterization protocols have been internationally accepted for TD diffuse optical systems: (1) the Basic Instrumental Performance (BIP) protocol, (2) the Optical Methods for Medical Diagnosis and Monitoring of Diseases (MEDPHOT) protocol, and (3) the Noninvasive Imaging of Brain Function and Disease by Pulsed Near Infrared Light (nEUROPt) protocol (*21*–*23*).

### BIP Protocol

The BIP protocol (Fig. 2A, B) evaluates basic characteristics of TD instrumentation which influence the quality and accuracy of measurements in clinical applications: responsivity of the detectors, differential non-linearity (DNL), afterpulsing ratio, system instrument response function (IRF) and system stability (Methods).

**Figure 2.**
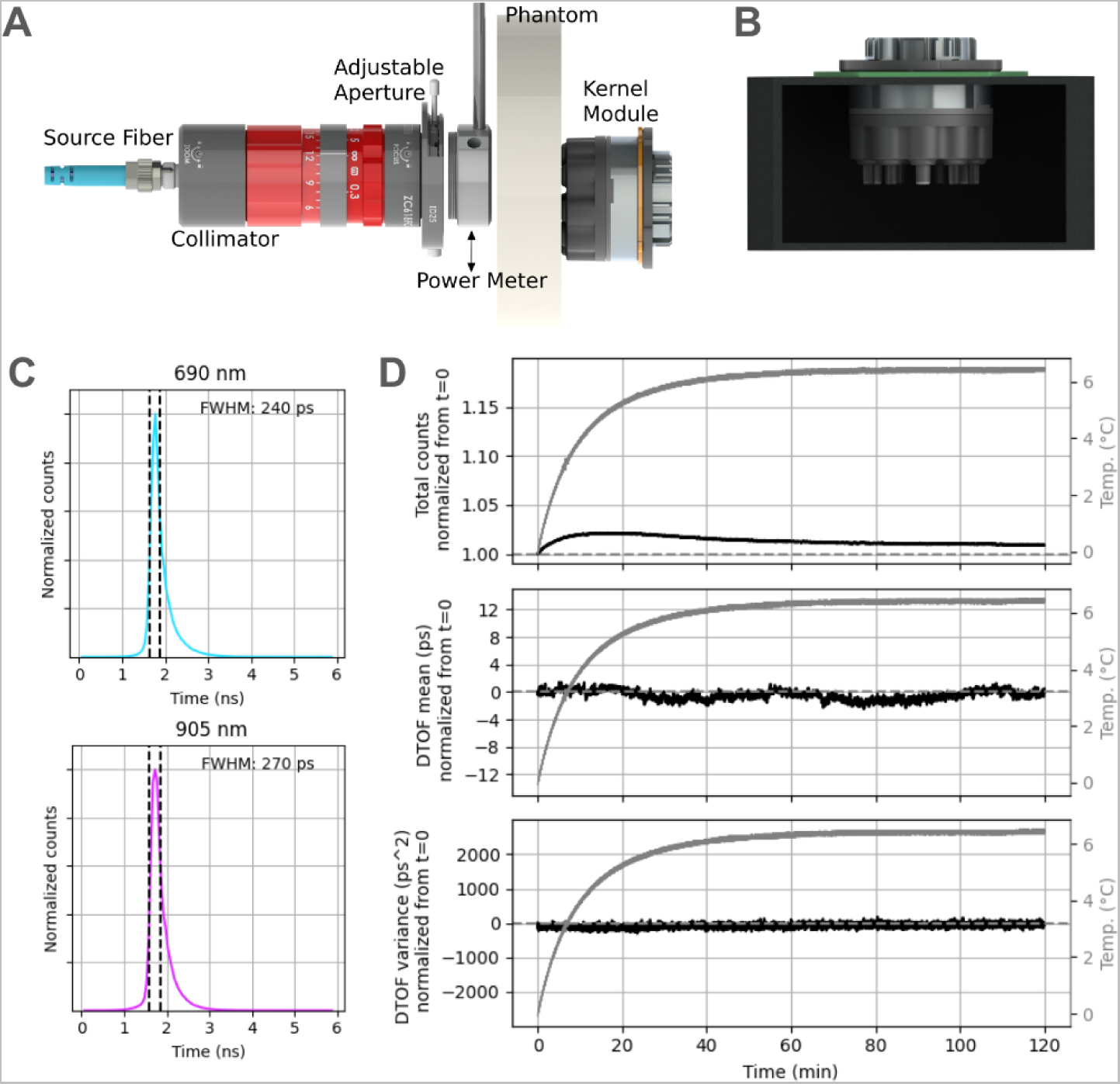
BIP protocol setup and results. **(A, B)** Schematic of the experimental setup for BIP experiments. (**C**) The continuously-recorded system IRF exhibited narrow widths for the two wavelengths (rows). Dashed vertical lines indicated the boundaries over which FWHM was computed. (**D**) Despite the change in temperature over a 2-hour-long recording session (gray, repeated on all plots), the three moments of DTOFs (rows, black) were stable when accounting for the IRF changes throughout the recording. Normalization for the three moments were done by division over the value at the first time point for the total counts, and by subtracting the value at the first time point for the mean and variance moments. Note that, although our system is capable of sampling much faster, an integration time of 1s was used for this demonstration, as is standard throughout the BIP protocol.

To assess the responsivity of the Flow2 detectors at the system’s two operating wavelengths (690 nm and 905 nm), we used an external laser to provide known input power (0.2mW). The corresponding responsivity was calculated to be 9.97x10^-9^ and 2.28x10^-9^ m^2^sr respectively at the two wavelengths. These are within the range of reported values in research-grade systems (*24*). The DNL was assessed under constant illumination (Methods; eq. 1), and was found to be 0.148. This DNL is relatively high compared to other TD devices (*24*). However, there isn’t an established threshold of acceptable DNL for TD systems, and we have not found the DNL of Flow2 to be detrimental to any of the functional characterization results (see next sections, MEDPHOT and nEUROPt).

Other metrics were assessed during normal operation of the system. We recorded a two-hour session of data using a single module, mounted on a custom fixture to capture light in reflectance mode (Method). We examined the data from the detectors that capture light that traveled externally (reflected on matte surface of the fixture), as well as from the module’s dedicated IRF detector (which receives light directly from the module’s lasers). The afterpulsing ratio (Methods; eq. 2), a known source of signal-dependent noise, was 0.0024 at 690 nm and 0.0019 at 905 nm. While this metric is not as widely reported in the literature despite being part of the benchmarking protocol, these values are similar to those reported for another system (*25*).

The IRF of a TD system is critical to its ability to accurately characterize the media through which light travels. The data recorded by the system for a given source-detector pair (which we refer to as the DTOF) consists of the convolution of the DTOF one expects from the radiative transfer equation (which we refer to as the temporal point spread function, or TPSF) with the IRF of the system (*26*). Therefore a wide IRF tends to blur information. For our system, the full-width at half maximum (FWHM) of the IRF (as measured on the IRF detector) was 240 ps at 690 nm, and 270 ps at 905 nm, at the beginning of the session (Fig. 2C). At the end of the 2-hour session, it was 285 ps at 690 nm and 305 ps at 905 nm. This drift with recording time is due to an increase in temperature over the course of the session, which affects several aspects of the electronics. The IRF FWHM is below or similar to the mean as compared to other systems (*24*). While the IRF is typically characterized by its FWHM, we note that the ability of a TD system to accurately characterize media with higher absorption is determined by the slope of the tail of the IRF. We therefore also report the width at other fractions of the maximum (and encourage others to do so). We found that the IRF width at 10% of the max was 710 ps at 690 nm and 710 ps at 905 nm at the beginning of the session, vs. 710 ps and 725 ps respectively after 2 hours. The width at 1% of the max was 1630 ps at 690 nm and 1655 ps at 905 nm, vs. 1740 ps and 1715 ps respectively after 2 hours.

A common way to summarize information from time-of-flight histograms is to compute the first three moments of the histogram corresponding to the total counts (sum), mean time-of-flight (first moment), and variance of the times of flight (second central moment) (*13, 27, 28*). Moments have a convenient property: the moments of the DTOF can be obtained from calculating the moments of the TPSF and of the IRF straightforwardly (*27*). Accordingly, with Flow2, system drift in the DTOF moments can be corrected for, using the internal IRF detector. We demonstrate here that the first three moments of DTOFs (total counts, mean, variance) were consistently stable over the course of the 2-hour session after such correction, as shown for a representative channel in Fig. 2D.

### MEDPHOT Protocol

Next, we probed the accuracy of Flow2 measurements in retrieving known optical properties of homogeneous phantoms using the MEDPHOT protocol. Being able to retrieve accurate optical properties at each wavelength used by the system is a prerequisite to being able to retrieve accurate estimates of the concentrations of oxy- and deoxyhemoglobin from the combination of the wavelengths (through the Modified Beer-Lambert Law). Here, a single module was used to characterize absorption and scattering coefficients (μ_a_ and μ_s_′, respectively) of 12 solid phantoms (BioPixS; cylindrical in shape; 50mm height, 100mm diameter), with varying absorption values (denoted with numbers 1, 3, 5 labels) and different scattering values (labeled by letters A-D) (Fig. 3A). Importantly, the range of properties represented by these phantoms encompasses properties of human head tissue (*29*) (skin, skull, cerebrospinal fluid, gray matter, and white matter) (Fig. 3B). Both μ_a_ and μ_s_′ were estimated using conventional curve fitting method: an iterative search using the Levenberg-Marquardt algorithm, seeking to minimize differences between measured DTOFs, and model DTOFs derived from convolving the measured IRF with analytical TPSFs (Methods).

**Figure 3.**
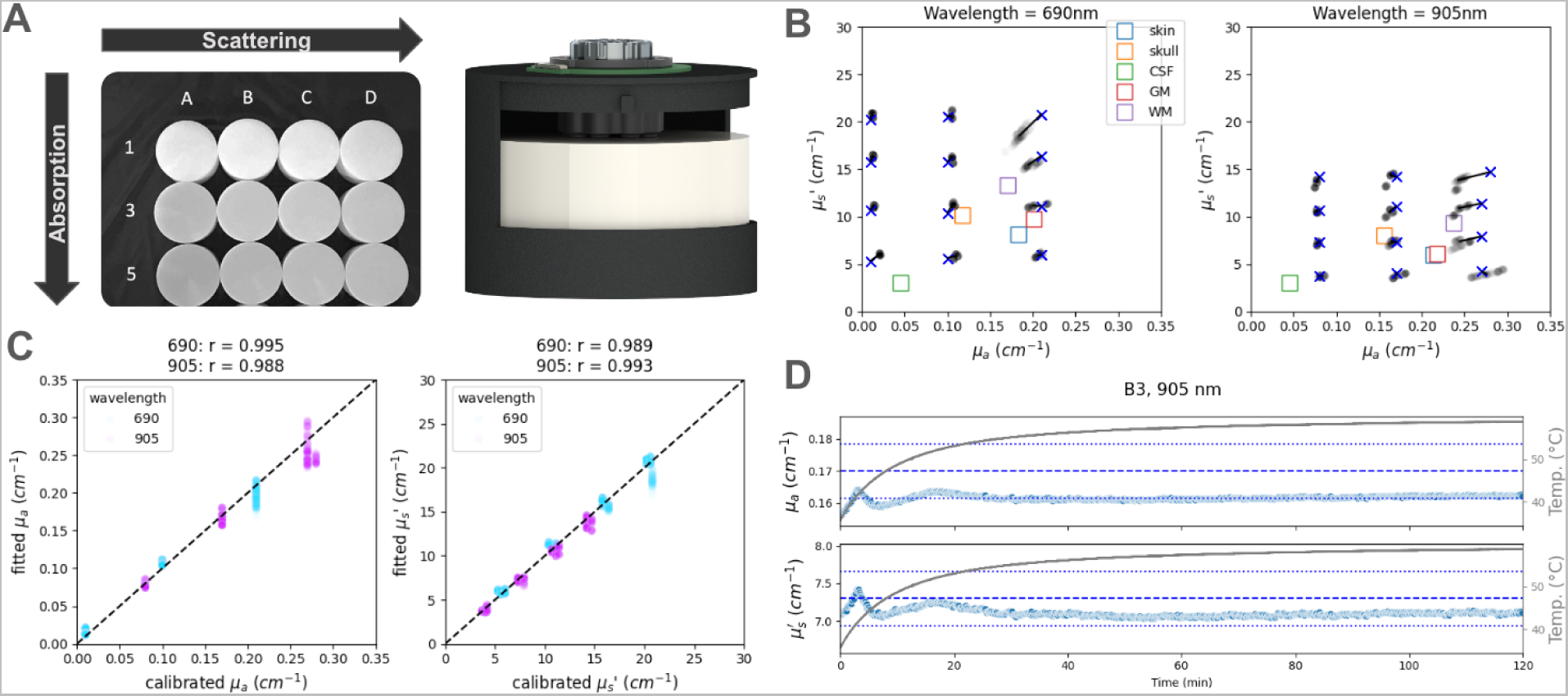
Retrieving known optical properties using the MEDPHOT protocol. **(A)** Twelve different phantoms with known absorption and scattering coefficients (left) were used in turn in our MEDPHOT recording setup (right). **(B)** Retrieved optical properties for all phantoms in the matrix (left: 690nm; right: 905nm). The blue x’s denote the calibrated values supplied by the phantom manufacturer. Each light black dot corresponds to the result from a single within module channel with source-detector distance 26.5mm, for a single 1s integration sample. The colored squares correspond to known average tissue properties in the human head, and are added for reference (CSF: cerebrospinal fluid; GM: gray matter; WM: white matter). **(C)** Correlation between retrieved and known values of μ_a_ (left) and μ_s_’ (right), for both wavelengths (cyan: 690nm, magenta: 905nm). **(D)** Stability of optical property retrieval over a two hour measurement, from cold start (temperature of the laser, displayed in gray, increases by 25 deg celsius during the session). The results are from a single channel, with 1s integration time. The dashed blue line indicates the value supplied by the manufacturer, and dotted blue lines indicate +/-5% from this value.

We found the retrieved optical properties to be very close to the calibrated values for both wavelengths and all 12 phantoms (Fig. 3B), with the following average relative errors: 8.4% and 4.3% for μ_a_, and 4.2% and 3.8% for μ_s_′ at 690 nm and 905 nm respectively. These relative errors are similar to what has been reported for other instruments (*24*). We note that accuracy is poorer at higher absorption levels, which we primarily attribute to a limitation imposed by the slope of the IRF (hence the importance of characterizing it fully). The retrieved and calibrated optical properties showed excellent linearity, with a correlation coefficient over 0.98 for both μ_a_ and μ_s_′ (Fig. 3C), and a median relative error with respect to the line of best fit of 6.3% (690 nm) and 3.0% (905 nm) for μ_a_, and 2.3% (690 nm) and 3.1% (905 nm) for μ_s_′. This performance is in the range of those reported by similar instruments (*24*).

We also tested the stability of the retrieved optical properties over a 2-hour recording period (for phantom B5). Despite a noticeable increase in the system temperature (Fig. 3D), the retrieved μ_a_ and μ_s_′ were stable with no noticeable trend (slopes computed over full time range are 0.005%/min for μ_a_ and -0.010%/min for μ_s_′) further confirming that the continuous measurement at the dedicated IRF detector is an effective way to fully correct for system drift.

### nEUROPt Protocol

The nEUROPt protocol is a system-level evaluation method that utilizes optical phantoms mimicking brain tissue (*22*). We used previously described methods to perform this evaluation (*14*) (Fig. 4A, Methods). Briefly, this procedure involves a liquid phantom (a mixture of water, India ink, and intralipid emulsion) titrated to have optical properties of μ_a_ = 0.01 and μ_s_′ = 1.0 mm^-1^ (prepared separately for each wavelength). Experiments were conducted using black polyvinyl chloride (PVC) cylinders of various sizes, suspended in the liquid phantom, and moved incrementally away from the source-detector plane. Measurements were taken at depths ranging from 8mm to 36mm, following the original protocol’s guidelines of 100 1-second accumulation histograms.

**Figure 4.**
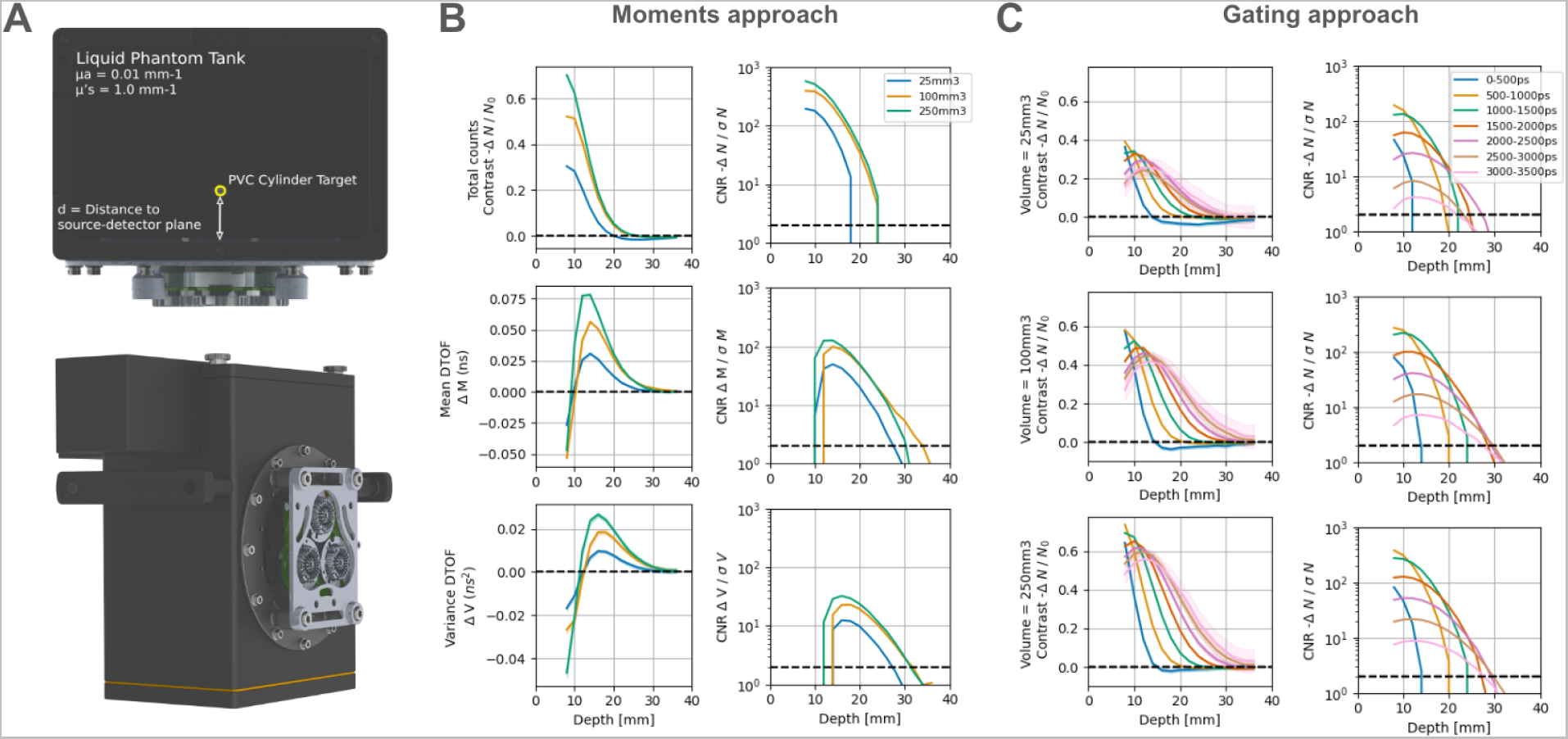
nEUROPt protocol setup and results. **(A)** Schematic of the setup used to test the nEUROPt protocol. **(B)** Left) Contrast as a function of depth for three different moments of DTOFs (sum, mean and variance are the three rows respectively). Each color represents the volume of a different PVC cylinder used in the experiment. The black dashed line emphasizes the position of zero contrast. Right) Same as (left) but the contrast-to-noise ratio (CNR) at different depths. The black dashed line shows a CNR of 2, which we arbitrarily chose as the limit above which a signal can be reliably detected. **(C)** Contrast (left) and CNR (right) are shown as a function of depth for the three different PVC cylinders used (rows). Each color corresponds to different time gates used for analysis. Both B and C are results with the 905 nm wavelength and a representative channel. For the results on the 690 nm wavelength, see Fig. S1. The results for both wavelengths, when using the time integration method yielded qualitatively similar results to the counts integration method (shown here).

As is customary in TD-fNIRS analysis, we present results after summarizing the full resolution DTOFs in two different ways (*13, 28, 30*): (1) using the first three moments, as described in the previous section; (2) using coarse time gates: here, we first deconvolve the measured IRF from the measured DTOFs, and coarsen the data to 500ps gates (0-500ps, 500-1000ps, etc). For each set of features (moments and gates) and each wavelength, we computed two different metrics: contrast, and contrast-to-noise ratio (CNR), as a function of occlusion depth (Methods).

For the moments, we found that both contrast and CNR metrics exhibited a depth-selective profile with higher moments being more sensitive to deeper occlusions (Fig. 4B), as expected from prior literature (*28*). Similarly, when using the gating approach (Fig. 4C), later time gates show better contrast and CNR with depth (*31*). It is important to note that with both methods, there is high contrast and CNR at depths > 20mm (which is well into the brain) (*32*), especially for higher moments and later time gates—demonstrating the advantage of TD-fNIRS systems over CW-fNIRS systems with respect to depth sensitivity. These results are among the best performance of TD systems according to a recent benchmarking study (*24*).

### In-vivo recordings of absolute brain oxygenation during a breath hold challenge

The breath hold task consisted of blocks of a transient hypercapnic challenge (a 20 second breath hold following an exhale), alternating with periods of visually-guided paced breathing (Methods). This is a commonly used calibration task in the hemodynamic neuroimaging literature, as it produces a large and reproducible response in blood oxygenation and blood flow in the brain (*33*).

To obtain dynamic estimates of the absolute concentrations of oxygenated and deoxygenated hemoglobin (HbO and HbR respectively), we used previously described methods (*14*–*16, 34*). The DTOFs recorded from the Flow2 system underwent standard preprocessing and cleaning steps. We then applied the curve fitting method (as in the MEDPHOT protocol), assuming a homogeneous medium. This is of course a simplification, as head tissue is not homogeneous. We observed systematic changes in the absolute HbO and HbR that were locked to the task events (Fig. 5A) and these increases and decreases were consistent with prior literature (*33, 35*). Specifically, HbO revealed a significant increase after the start of the breath-hold period and underwent a decrease when the next breathing period began. Furthermore, both HbO and HbR were modulated by the breathing pattern such that inhaling and exhaling resulted in changes in the signals (inhale: HbO increases, HbR decreases; exhale: opposite). Lastly, we extracted the time-varying heart rate (using the dedicated prefrontal module; Methods), which revealed the presence of strong respiratory sinus arrhythmia (RSA; Fig. 5B), i.e., modulation of the heart rate as a function of the phase of the breathing cycle (*36*).

**Figure 5.**
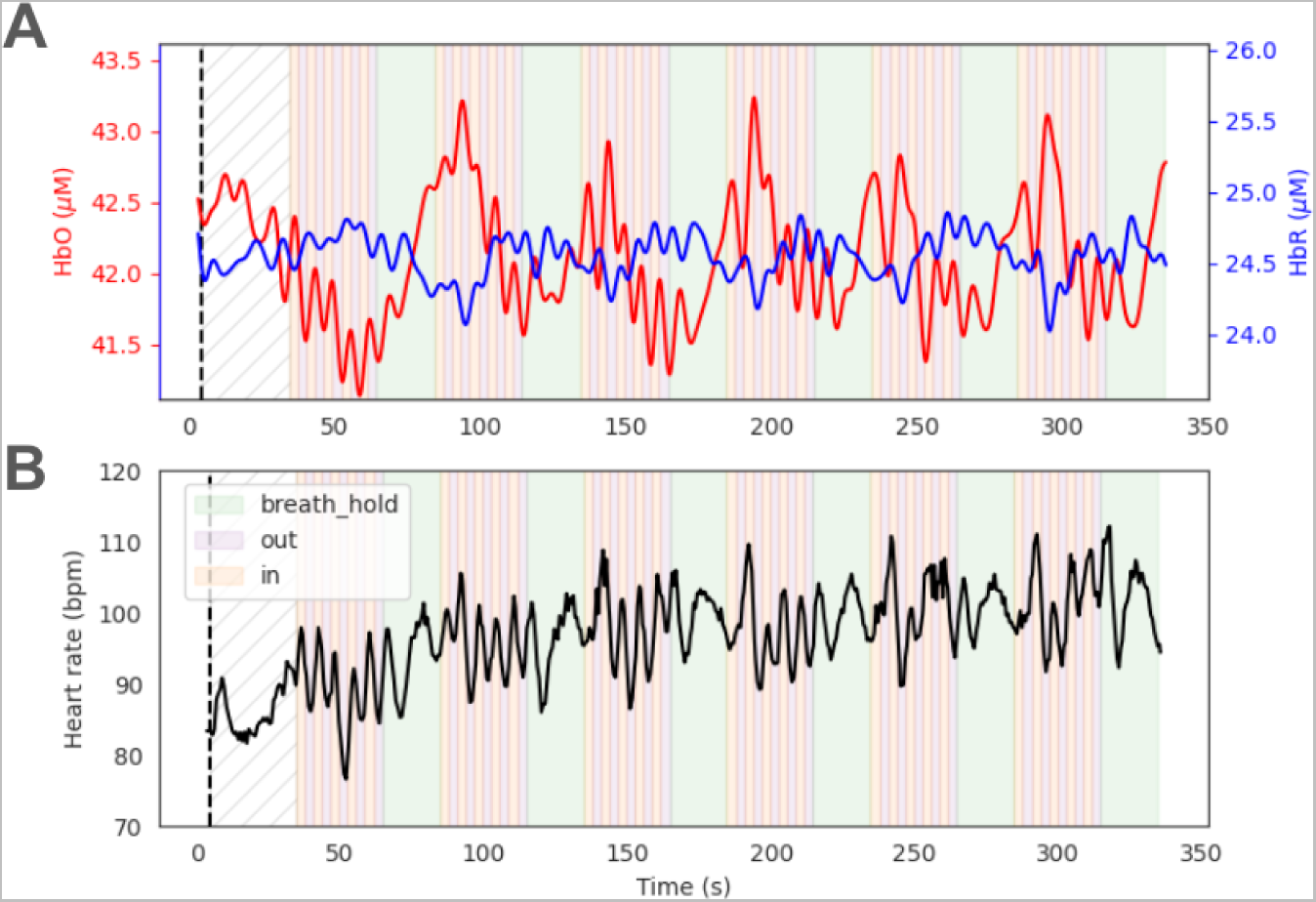
Absolute changes in the concentrations of oxygenated and deoxygenated hemoglobin in a breath hold task. **(A)** Retrieved absolute concentrations of HbO (red) and HbR (blue) were separately averaged over all long within-module channels for two well-coupled prefrontal modules. Note the gradual increase (decrease) in HbO (HbR) during the breath holding portion of the task. During rhythmic breathing, the signals reflected the pattern of breathing (faster oscillation during in/out periods). Periods of breath holding and breathing (inhale/exhale) are shown with different colored backgrounds. **(B)** Using a dedicated fast-sampling prefrontal module, we were able to extract the heart rate time course during the experiment. The modulation of heart rate with breathing (RSA) was visible in the data (higher frequency oscillations during the breathing phase).

### In-vivo recordings of brain oxygenation changes during a sensory and a motor task

In order to validate Flow2’s ability to record robust sensory-related brain activity, we employed a passive auditory task. During this task, the participant wore earbuds and was asked to listen to clips of brown noise (noise blocks) and clips of human speech (story blocks) that were interleaved in a pseudorandomized manner. We utilized a standard Generalized Linear Model (GLM) approach that consisted of modeling the hemodynamic signal recorded during the task (for each channel) as a function of the experimental design conditions (story and noise blocks) (Methods). This analysis revealed significantly more activity in bilateral auditory cortex during the story blocks than during the noise blocks (Fig. 6A)—a pattern of brain activation consistent with the literature (*37, 38*).

**Figure 6.**
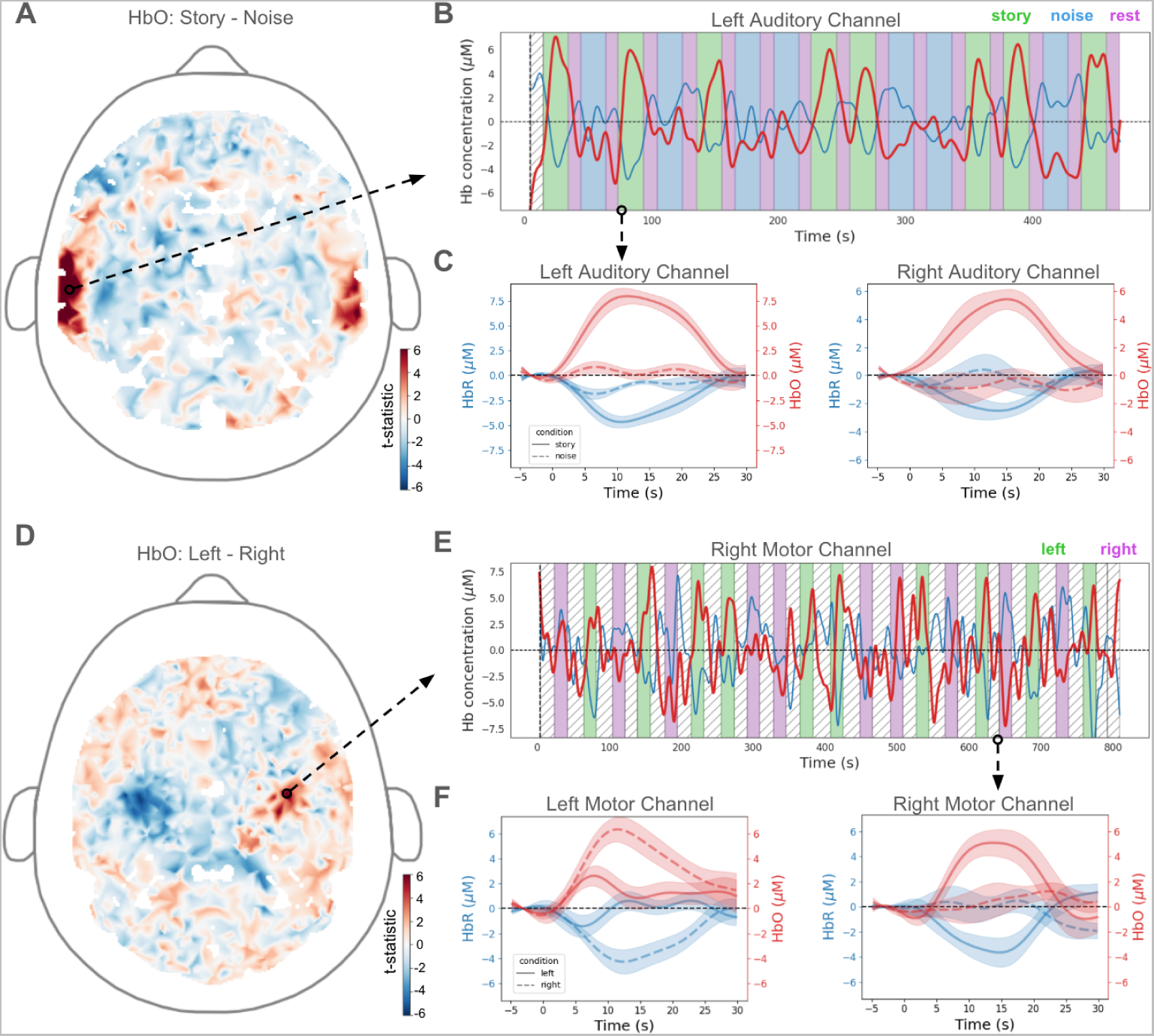
Task-related hemodynamic activity extracted from Flow2 human recordings. **(A)** GLM-derived pattern of brain activity (channel-level test statistics for the story-noise contrast) over the whole-head showed regions of significant activation in bilateral auditory cortex (warmer colors). Arrow indicates the regions where the representative channel, shown in (**B**) is located. **(B)** Full time course of HbO (red)/HbR (blue) for a representative channel from left auditory cortex overlaid on the task design (green: story blocks, blue: noise blocks, purple: rest periods). **(C)** Averaged epoched responses (mean ± standard error) for representative channels from left/right auditory cortex (left/right panels, respectively). Left Auditory is the same representative channel as (**B**), as indicated by the arrow. Note the increase in HbO (red) and decrease in HbR (blue) during the story condition (solid lines), as well as the muted activation/deactivation to the noise condition (dashed line). **(D)** Same as (**A**), but for the left - right contrast during the finger tapping task. Notice regions of significant activation (warmer colors) and deactivation (cooler colors) in the right and left motor cortex, respectively. **(E)** Same as (**B**), but for a representative channel from the right motor cortex, as indicated by the arrow. Notice the increase in HbO during the left finger tapping blocks (green epochs) compared to the right finger tapping blocks (purple epochs). **(F)** Epoched responses as in (C), but for a channel within the left and right motor cortex. Notice the reversal in activation by condition across the hemispheres, where the left motor channel shows an increase in HbO to right finger tapping and the right motor channel shows an increase in HbO to left finger tapping.

To confirm the statistical output of the GLM visually, we examined the channel-level oscillations of HbO/HbR concentration over the full time course of the task. When overlaid with the block design of the task, channels that were within the regions of interest (and identified as significant by the GLM analysis) demonstrated the expected hemodynamic activity, effectively tracking the experimental blocks throughout time (representative channel from left auditory cortex shown in Fig. 6B). As expected, the two chromophores fluctuated in opposing directions.

Second, for each channel, we averaged the hemodynamic signal over epochs corresponding to the same block type. Representative epoched responses for single channels located in the right/left auditory cortex are displayed in Fig. 6C. These channels exhibited increases in HbO, and corresponding decreases in HbR during the story condition of the task. Notably, they also exhibited negligible or relatively smaller responses to the noise condition and complementary HbO/HbR activity.

As a further validation of the system, we employed an active motor task and repeated the exact same analysis (as we did for the passive sensory task). Specifically, we recorded the brain activity of a participant engaged in a classical finger tapping task, during which they were repeatedly cued to use either their left or right hand and tap their fingers sequentially to their thumb for a period of time (Methods).

Again, we first validated the whole-head brain activation patterns via a GLM approach. Unlike auditory stimulation, which exhibited coherence in temporal activation across hemispheres, here we expected the motor cortex to demonstrate opposing activation across hemispheres to task conditions (*39*). Indeed, this analysis revealed regions of significant activation in the right motor cortex and regions of significant deactivation in the left motor cortex when comparing left-hand tapping to right-hand tapping (Fig. 6D).

The time course for a representative channel from the right motor cortex exhibited increases in HbO throughout time that were matched to left blocks and accompanied by decreases in HbR (Fig. 6E). These characteristic hemodynamic responses were even more evident after epoching. The epoch-averaged time courses for representative channels exhibited condition-specific hemodynamic responses. Specifically, a channel in the left motor cortex showed a rise in HbO and a corresponding decrease in HbR during right-tapping blocks (dashed line) (Fig. 6F; left), while a channel in the right motor cortex showed increases and decreases in HbO and HbR, respectively, during left-tapping blocks (Fig. 6F; right). Taken together these analyses underscore the ability to use Flow2 to record meaningful brain activity during a passive sensory task as well as an active motor task.

### Reconstructing brain activity with Time Domain diffuse optical tomography

High-density fNIRS recordings enable tomographic reconstruction of brain activity; recently, HD-DOT has been validated against fMRI for mapping the functional dynamics of the human cortex (*29*). To date, HD-DOT for the whole human head has only used CW-fNIRS. Given the advantages afforded by TD-fNIRS, notably in terms of depth sensitivity, we hypothesized that our system could yield better tomographic reconstructions than are possible with CW data alone. We used the datasets discussed above for passive auditory and finger tapping tasks to test our hypothesis.

It has been shown that different moments of the DTOFs exhibit different depth sensitivities (Fig. 4B, also see (*28*)). To assess how the quality of reconstruction varies when using only the sum moment (i.e., intensity—similar to what CW systems measure) versus when additional moments (mean and variance) enabled by TD capabilities are included, we obtained reconstructed HbO/HbR cortical maps in these two scenarios (Methods). The time course of reconstructed activity was further analyzed using GLM, which resulted in statistical maps for each task; specifically for Story-Noise contrast during the passive auditory task, and Left-Right tapping during the finger tapping task (Methods).

Activation maps from the sum moment exhibited expected patterns for the auditory task: increased activity during story blocks as compared to noise blocks in the auditory region (Fig. 7A). In the finger tapping task, too, the activation maps revealed elevated contralateral motor cortex activity (Fig. 7B).

**Figure 7.**
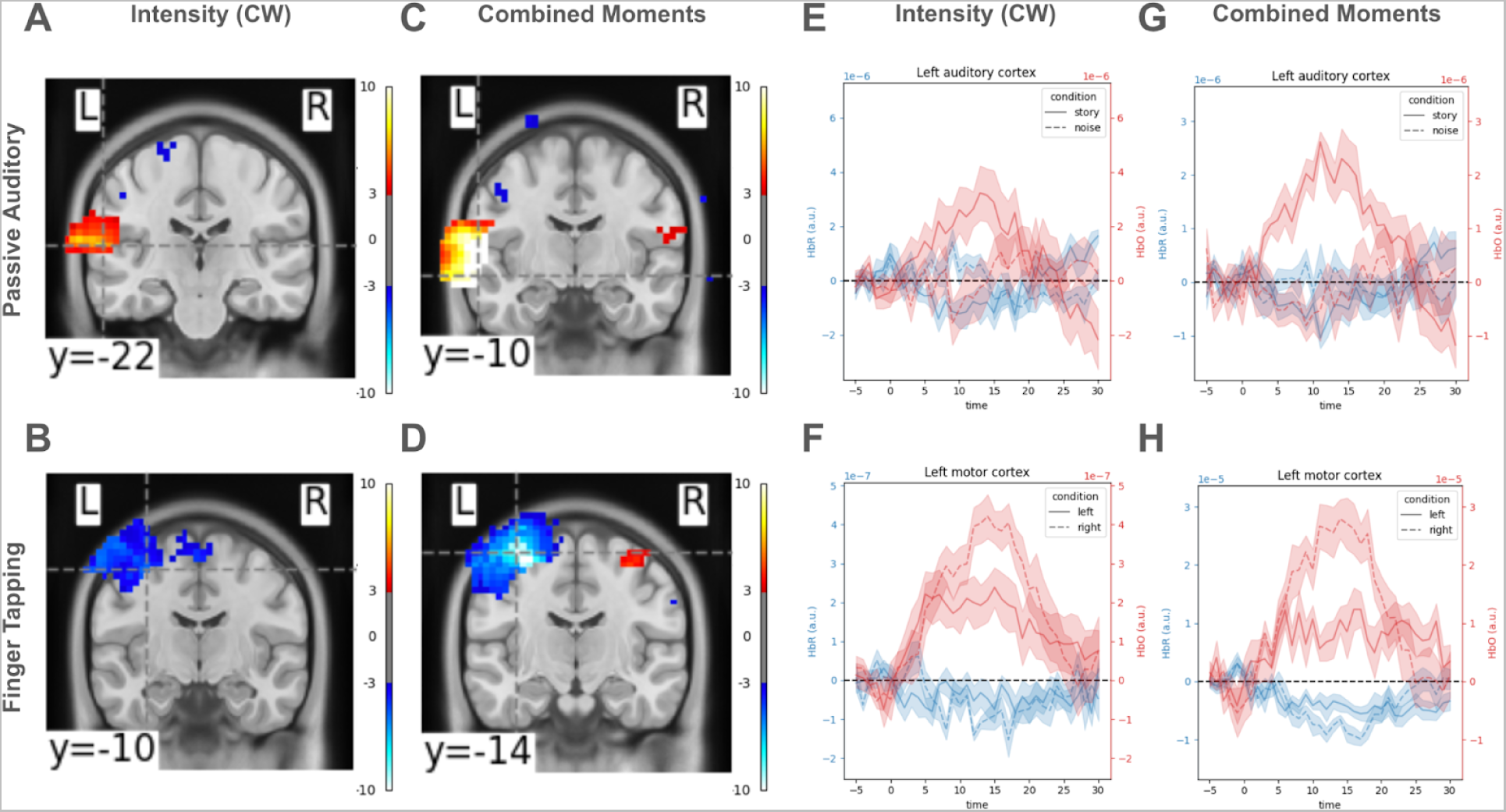
Cortical activations maps reconstructed from a sensory and a motor task. **(A)** GLM-derived statistical map (t-statistic) of brain activation for Story - Noise contrast revealed an area with high activation in the left auditory cortex when using only the intensity data to reconstruct HbO/HbR. **(B)** Same as (A) but results are from the finger tapping task for the Left - Right tapping contrast. **(C, D)** Same as (A, B) but when reconstruction was done using all three TD moments (i.e., sum, mean, and variance). Note the larger values of t-statistics in these statistical maps. **(E)** Stimulus-locked average time course of voxels within a 10 mm radius spherical seed centered at the brain voxel with the highest activation in the left auditory cortex (dashed gray crosshair in (A)), when using only the sum moment to reconstruct brain activity. **(F)** same as (E) but results are from the finger tapping task, and the brain voxel with highest contrast is denoted by the crosshair in (B)**. (G, H)** Same as (E, F) but when reconstruction was done using all three moments. Note that, following the original downsampling to 1Hz, the time courses have not undergone any further smoothing or filtering. Also, note that the y-axes have different scales, to best visualize the dynamic range of the data for each independent reconstruction.

When performing reconstruction using all three TD moments, we indeed found deeper sensitivities with higher moments (Fig. S2). Consequently, both tasks demonstrated even larger, more significant, statistical contrasts between activations during the conditions in expected regions (Fig. 7C, D). This difference was even more stark when considering the time course of activation in predetermined task-specific ROIs when they were time-locked and epoched (Fig. 7E-H). For example, while the sum moment-based HbO time course in the left motor region during right-tapping blocks exhibited small elevated activity compared to that during left-tapping blocks (Fig. 7F), this contrast in activity was notably larger when using the reconstructed HbO from all moments (Fig. 7H).

## Discussion

In the current study we introduce and validate the Kernel Flow2 as a wearable TD-DOT system capable of measuring and reconstructing brain activity over the whole head. We have shown the system has performance comparable to limited channel count research-grade devices according to standardized benchmarking protocols, while also extending the field-of-view over the whole head. The system hardware was developed using scalable methods used for consumer electronics such as custom ASICs, plastic molded optics, low-power application processors, and standard communication protocols. This combination of system performance, cost, and form factor can enable many applications for both neuroscience research and precision neuromedicine.

It is critical to validate the performance of newly developed measurement systems using standardized protocols, before they can be applied for research or clinical purposes. For TD-fNIRS systems specifically, recent multi-laboratory efforts have resulted in comprehensive tests to characterize performance: the BIP, MEDPHOT, and nEUROPt protocols. Each of these focuses on performance from a different angle (*21*–*24*). Briefly, these tests revealed the following for Flow2: (1) the system’s IRF is adequately narrow and stable, which is required for various computations (e.g. computing absolute HbO/HbR concentrations); (2) the system can indeed be used to accurately retrieve optical properties from calibrated optical phantoms with optical properties in the physiological range at both wavelengths, which is required to retrieve accurate concentrations of HbO and HbR; (3) the measurements from the system have better depth sensitivity than the measurements from CW systems (intensity only). These metrics were similar or better than those reported for limited channel count systems (*24*). Together, these validations against known ground truth suggest that Flow2 recordings should be of high enough quality to study hemodynamic activity on a human head. We thus proceeded to investigate spatial and temporal patterns of brain activity extracted from neural recordings in vivo on a human subject.

First, we used an easily implemented mild hypercapnic challenge, which consists in a short breath hold (at the end of an exhale). This challenge is known to trigger an increase in arterial PCO_2_, resulting in a homeostatic response including cerebral vasodilation and an increase in cerebral blood flow; importantly, this mild challenge does not appear to result in extracranial perfusion effects (*35*). We were able to use curve fitting with the TD data to track the absolute concentrations of oxygenated and deoxygenated hemoglobin, something that traditional CW systems cannot do without making additional assumptions. Because the homeostatic response is limited to cerebral tissue in this hypercapnic challenge, we can demonstrate that our approach of using a homogeneous model is sensitive to absolute oxygenation changes in the brain. Retrieving absolute properties is of particular interest in many clinical applications of TD-fNIRS such as in stroke assessment (*40*), neurocritical care (*41, 42*), multiple sclerosis (*43*), schizophrenia (*44*), and dementia (*45*). Thus far, because of the high cost and bulkiness of these systems, their clinical use cases have been limited to research settings (*30*). Therefore, using a portable system such as Flow2 may present an opportunity to further translate these lines of clinical research into practice.

We then had participants wearing the Flow2 headset engage in two simple tasks: a passive auditory (sensory) task, and a finger tapping (motor) task. Note that both of these tasks are classic neuroimaging paradigms with consensus from the scientific community regarding the foci of functional brain activity (*37, 38, 46*). Our analyses showcased localized regions of activation in agreement with prior literature: bilateral auditory cortex activity for the passive auditory task; and contralateral motor cortex activity for the finger tapping task. Further, those channels that exhibited the most statistically significant responses to task conditions revealed strong task-locked modulations. It is worth noting that we previously demonstrated the stability of such task-evoked metrics in a larger-scale study with the Flow2 device (*34*). Taken together, these findings uphold the two pillars of device performance for research and clinical-use: validity and reliability.

Individual channels of the Flow2 system yield high quality hemodynamic signals from the human head. As the Flow2 system is composed of a very large number of channels (over 3500 with source-detector distance < 60mm) in a wearable helmet-like form factor, it further enables reconstruction of brain activity (DOT) over the whole head. The density of channels is on par with state-of-the-art CW HD-DOT systems (*47*). Furthermore, TD-DOT improved the statistical significance of cortical activations as compared to CW-DOT; we can speculate that this is because of its better depth resolution as demonstrated in our characterization results (e.g., nEUROPt protocol results). We note that the comparison performed here is as targeted and unconfounded as can be, as it was performed using the same input data (TD DTOFs) summarized in different ways. Establishing that in vivo reconstructed activations have better anatomical specificity in the TD-DOT case than in the CW-DOT case would require an independent measurement, e.g., using fMRI, to establish a ground truth to which reconstructions could be compared.

While the Flow2 system has overcome many challenges of the previous prototype Flow1 system and low-channel-count TD systems, there are still remaining limitations. The full headset remains relatively heavy, as all electronics are built-in. The weight may limit system wear time, thus the headset may not accommodate applications requiring long recordings (although, in some cases this can be mitigated with thoughtful study design). Additionally, as with other optical neuroimaging devices, hair can limit signal quality, as it absorbs light which reduces signal at the detectors. We have found that at least 1000 usable channels are available over the head for most participants (*34*) there is still room for improvement in mechanical design to ensure good optical coupling by working through all hair types and making contact with the scalp at all locations. Moreover, as with all diffuse optical systems, deep brain structures remain inaccessible. The TD nature of our device does however allow for deeper sensitivity in cortical regions as compared to CW systems. Finally, to perform a reconstruction of brain activity, a structural head model must be used which describes the tissue composition (scalp, skull, cerebrospinal fluid, gray matter, white matter) and the precise positioning of all optodes (sources and detectors) must be known. For the reconstructions performed in this work, we used an atlas head model and average optode positions (see Methods). This is the most scalable approach, which does not require individual head measurements or individual structural MRIs, and solely relies on good practices for headset positioning. It is also an approach that makes several assumptions and therefore has room for error. There are intermediate ways to improve atlas-based tomography by using individual measurements, such as individually registering optode locations (*48*) or picking the best atlas from a library (*49*). There are ongoing efforts to make such methods readily available for Flow2. Additionally, more advanced tomographic reconstruction approaches that fully utilize the DTOFs will be explored and implemented in future iterations (*50*).

Although the application of optical neuroimaging systems in clinical settings has been discussed repeatedly (*8, 30, 51, 52*), we argue that no system to date has successfully integrated the combination of: 1) an easy-to-use form factor; 2) the most advanced fNIRS modality (TD); and 3) high enough density to enable the reconstruction of whole-brain cortical hemodynamic signals. The confluence of these factors in Flow2 facilitate better, larger-scale, and more timely data collection efforts. The combination distinctly positions the device as a premier recording option for both the scientific and clinical communities. Future studies with TD-DOT can translate insights gleaned from neuroimaging research and bring them into practice, therefore enabling novel advances in precision medicine through scalable neuroimaging-based biomarker identification (*53*–*55*).

## Materials and Methods

### Detailed description of the Kernel Flow2 system

#### System specifications

The system consists of 40 modules that are arranged in a headset design (Fig. 1A, B). Modules are organized into 7 headset superstructure plates that cover the frontal, parietal, temporal, and occipital cortices. Each Kernel Flow2 optical module consists of 3 dual-wavelength sources and 6 detectors located on a 13.5mm radius from module center. An additional detector located in the center of the module continuously measures the IRF. The 3 source emission points are offset 120 degrees, with a detector located 37 degrees on either side of each source point (Fig. 1C).

Intra-module channel distances are 8.5mm (6 source-detector pairs), 17.9mm (6 source-detector pairs), and 26.5mm (6 source-detector pairs), for a total of 18 dual-wavelength channels within a module. Cross-module channels also provide data, for a total of 2,565 possible channels with a source-detector separation of ≤ 50 mm over the whole head (3,583 ≤ 60mm). The actual number of usable channels depends on the light attenuation for a particular participant. Module locations are fixed within a plate, and plates are held together with an adjustable tensioning system. The overall size of the headset fits a range of adult head sizes, with a minimum size of 52-cm circumference and a 32.5-cm bitragion coronal arc. The system weighs 2.4kg. By comparison, a single detector channel wearable system was reported to weigh 2.5kg (*56*), although that system includes the weight of the battery while the Kernel Flow2 system is powered over universal serial bus type-C (USB-C) using the USB power delivery (USB-PD) standard.

Each module in the system consists of three major subassemblies: laser assembly, detector assembly, and the optical assembly. All three of these subassemblies are shown together in an exploded view in Fig. 1D. The details of each subassembly and the overall system architecture are presented in the following sections.

#### System architecture

The Flow2 system has a hierarchical architecture with electronics and wiring harnesses integrated into the headset. The system is cabled over a single USB-C interface that both supplies power (USB-PD) and enables bidirectional communication (USB 2.0) between the data collection computer and the Flow2 system.

The USB-C cable connects to the Flow2 system through the hub sub-assembly that includes an application processor (AP), and a global reference clock. In addition, the hub sub-assembly integrates 4 electroencephalography (EEG) analog-to-digital converter (ADC) channels that are designed for connecting to active dry electrodes. The hub also handles the primary power negotiation for the USB-PD standard and distribution of power to the rest of the system. Connected to the hub are follower boards that serve as data aggregation points for clusters of modules. Each of these follower boards include a low-power field-programmable gate array (FPGA), additional power conditioning circuitry, a 9-axis IMU, and local USB 2.0 communication interface between the follower and hub.

In total, the Flow2 system supports connection of up to 40 time-domain optical modules and includes 4 active dry EEG channels (approximate locations from 10-10 montage: AF4, AF3, FCz, CPz). The system may be operated with fewer than 40 optical modules, allowing for the removal of unnecessary modules to reduce headset weight or cost. In the current study, we opted to utilize the complete headset configuration to depict activation and deactivation regions across the entire brain during different tasks as described in the following sections.

#### Laser source design

Each module source location (3 per module) has two pulsed edge emitting laser diodes. The paired laser diodes are placed diametrically opposite each other and emit light into a combined micro-prism and lens optical element. The lasers are driven by custom-designed pulse shaping circuitry that efficiently generates laser pulse widths that are <= ∼225 ps (690 nm has a pulse width of less than 205 ps, and the 905 nm laser has a pulse width of less than 250 ps). The lasers operate in gain-switched mode, which enables the production of optical pulses that are shorter than the electrical pulse that drives them. In addition to the laser driver circuitry and laser diodes, this sub-assembly contains the power conditioning circuits for the module, as well as the connectivity of the optical module to the follower board which serves as a data aggregation point for clusters of modules.

#### Detector assembly design

The detector sub-assembly comprises seven detector ASICs custom-designed by Kernel, specifically optimized for conducting time-of-flight measurements in diffuse optical tomography. Notably, the Kernel Flow2 ASICs feature integrated time-to-digital (TDC) circuitry with the photodiodes, ensuring a seamless fusion of key functionalities. The detector ASICs have been engineered to accommodate high photon count rates exceeding 5 Gcps, demonstrating resilience against pile-up distortion.

A novel custom band-pass coating has been developed and applied to the detector ASIC package. This specialized coating serves as a selective filter, tailored to discriminate against undesired wavelengths within the spectrum, thereby optimizing the detection of photons within the desired spectrum wavelengths of the Flow2 lasers. The implementation of this coating helps to address the challenges associated with ambient light and extraneous signals, ensuring a maximal signal-to-noise ratio and heightened accuracy in TD-fNIRS measurements.

Photon arrival times, derived from on-chip TDCs, are systematically aggregated into histograms and communicated via a serial peripheral interface (SPI) bus to a dedicated FPGA. The synchronization of ASICs across all modules to a 20-MHz global reference clock facilitates the recording of temporally aligned signals between any source and any detector in the system. Furthermore, each detector ASIC incorporates dedicated high voltage bias circuitry, optimizing the bias for each detector within the system for stable operation at any temperature. Additionally, new circuit architectures were used to reduce power consumption while improving overall TDC performance compared to the Flow1 design.

Within each module, one dedicated detector at the center of the module assumes the role of an on-board IRF detector. This detector captures light, transmitted via a waveguide from the source optical path, yielding a per-pulse waveform that temporally corresponds to each pulse from each respective wavelength. The programmable integration time for constructing histograms on each detector spans from 1ms to 800ms, with the current configuration set at 3.5ms. Each histogram collected contains signals from only one wavelength. This means our histogram sampling rate is 285.7 Hz, and considering both wavelengths, the system is able to complete spectroscopic measurements at a rate of 142.9 Hz. To avoid optical crosstalk, all lasers are not enabled at the same time. This temporal multiplexing enables lasers in an 38-state pattern, completing a full cycle of data collection for all modules and wavelengths every 76 histograms, corresponding to a system sampling frequency of 3.76 Hz. In the full headset configuration, one source operates at 7.52 Hz, which is a frequency fast enough to accurately capture pulse rate, a complementary measure to the optical properties. This temporal multiplexing provides the additional benefit of bringing the average power per source down to a level that classifies as a class 1 laser device according to the United States Food and Drug Administration Federal Laser Product Performance Standard Code of Federal Regulations Title 21 Section 1040.10 (US FDA FLPPS 21CFR1040.10).

#### Module optics design

The module optics (Fig. 1D,E) have been carefully designed for several purposes. The optics conduct laser light from the laser diode sources into the scalp, couple light returning from the scalp to the detectors, conform to the curvature of the head, reduce interference from hair by parting it, isolate detected signal to a single detector, and maintain both source output power and detector measurement intensity independent of compression.

There are a total of 9 spring-loaded light pipes contained within the optical module, with 3 source light pipes having an optical exit area of 1.3mm x 1.3mm, and the 6 detector light pipes each with a 3mm diameter. Each of these 9 light pipes are optically isolated from one another in separate cavities to prevent optical cross-talk and signal contamination. Both the sources and detectors have a 2 lens optical system that maintains optical intensity throughout the compression range of the light pipe springs.

Per each source, there are two laser diodes that are placed offset from each source optic axis and directed into a custom built optical assembly. The light from the diodes immediately enters the source optic and is redirected by means of an integrated micro-prism with reflective silver coating. The light continues diverging within the first polymethyl methacrylate (PMMA) optical component until it hits the first aspheric surface.

A secondary source optic is a solid body optical component, also made out of PMMA, and is spring-loaded with 5mm of travel. The secondary source optic collects the collimated light from the first aspheric optic, and homogenizes the light prior to exiting the final surface of the light pipe and passing into the user’s scalp. This secondary source optic is also shrouded with an opaque covering to prevent light from leaking from a source directly into a detector light pipe outside of the module housing.

The receiving optics are also designed to use springs to comfortably conform to the user’s head. The input aperture of the detector light pipes is 3mm, with a rounded edge on the external interface for additional comfort. We have created another two lens imaging system, consisting of one plano surface, and 3 aspheric fresnel surfaces, in order to keep the received optical intensity constant at the detector, regardless of spring compression.

#### Continuous instrument response function monitor

A critical performance metric of a time-domain optical measurement system is the IRF. The IRF is a measure of the uncertainty associated with each timestamp recorded and is reflected in the histogram of accumulated events, when used in time-correlated single-photon counting (TCSPC) applications, as a smearing or broadening of the desired signal being measured.

Each component in the system plays a pivotal role in shaping the system’s IRF, with the laser and detectors emerging as the predominant influencers. The IRF is primarily shaped by the design specifics of the detector process, the laser driver circuitry, and the laser itself. Moreover, the IRF characteristics of these active components exhibit dependencies on the temperature and operating voltages of the electronics and optoelectronics, factors subject to temporal variations and fluctuations during a measurement.

A crucial element in the updated Flow2 module design is the incorporation of a dedicated reference IRF detector within each module (Fig. 1F). This detector is the exact same design as detectors employed in measuring signals from the scalp but is strategically isolated to capture light directly emitted from the lasers without traveling through tissue. This isolation provides a reliable estimate for both the IRF contribution from the detector (owing to its identical design and similar process as the signal-measuring detectors) and the IRF contribution originating from the laser. This IRF measurement is recorded continuously, at the same rate as that of the other detectors.

### Characterization protocols used for Time Domain instrumentation validation

#### BIP Protocol

The BIP protocol is designed to assess basic hardware performance of time-domain instruments such as detector responsivity, differential non-linearity (DNL), afterpulsing, the system IRF, and system stability (*21*). The protocol is applicable to instruments based on pulsed laser sources with repetition rates of the order of several tens of MHz, fast single-photon detectors, and time-correlated single photon counting (TCSPC).

The detector responsivity metric in the BIP protocol measures the relative sensitivity of light detection for the system, calculated as the ratio of measured photons exiting from a calibrated phantom to the input illumination (*21*). The calibrated phantom is a diffuse medium, to ensure that the measurement accounts for collection efficiency of the optics as well as the detection efficiency of the detector. The experimental setup consists of a calibrated phantom illuminated on one side with a collimated laser beam emitted from an externally calibrated fiber source and coupled into a lens collimation unit to produce an approximately pencil-sized laser beam and measured on the other side with a Flow2 module detector as shown in Fig. 2. A power meter was placed between the collimated beam and the input face of the phantom to verify the correct power reading prior to each measurement and data collection session. We used an input power of 0.2mW to yield target counts similar to those reported in the BIP protocol (we reached 0.68x10^6^ photons and 1.12x10^6^ photons at 690 nm and 905 nm respectively, with our system’s default integration time of 3.5 ms).

The DNL measurement in the BIP protocol characterizes variability in the time bin width of the TDCs. Variability in the time bin width results in a non-uniform number of photons in each bin when the TDC is illuminated with a continuous light source. The DNL was measured by applying a uniform illumination source to the detector from a battery-powered CW light source for 100 seconds, and calculating the relative differences in the number of collected photons per bin. The deviation from the ideal of an equal number of photons in every bin is calculated as the peak-to-peak difference between the maximum and minimum bins, normalized by the mean photon counts over bins:

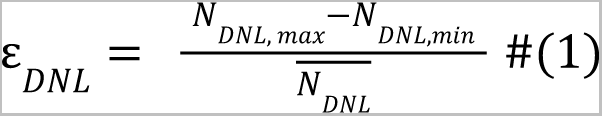

The IRF is carefully characterized in the BIP protocol to describe the overall time resolution of the system. It is typically measured by coupling the attenuated output of a source directly into a detector. The IRF was measured both at a typical detector and at the dedicated IRF detector. To measure the IRF at a typical detector and because our system doesn’t have the same flexibility of fiber-based systems, a custom fixture was used to capture light in reflectance mode. The source beam was adjusted to avoid saturating the detectors with direct illumination, and the light is then reflected off a matte surface to redirect it into the collection optics. To measure the IRF at the dedicated IRF detector, the built-in wave-guide that directly couples light from the source optical train to the IRF detection ASIC was used.

The IRF results from a convolution of the laser pulse shape and the temporal response of the detector and associated electronics. Per the protocol (*21*), the IRF was measured by averaging 20 histograms of 1s acquisition time. Because our detector maximum integration time is only 800ms, we collected these 20 histograms by summing individual 3.5ms histograms to produce a single 1001ms histogram. These 20 summed histograms were then averaged to calculate the IRF. The BIP protocol specifies a count rate of 1e6/s, i.e. each of the 20 1-s histograms should contain 1e6 photons. As the count rate of our system is much higher than 1e6/s, we also present the results when we match the protocol in terms of photon counts (i.e., much shorter integration time).

From the IRF measurement, we also calculate the afterpulse ratio (RAP), a signal-intensity noise source associated with the detector, as defined in the BIP protocol:

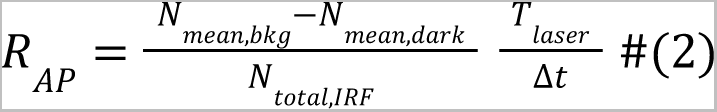

where N_mean,bkg_ and N_mean,dark_ are the average counts of the background measurement in the tail of the IRF and dark count measurements respectively, T_laser_ is the full laser period (1 / repetition rate) and dt is the time bin width. The afterpulsing ratio was calculated for both 690nm and 905nm. The afterpulse ratio is a measure of the increase in the noise floor due to this intensity-dependent noise.

Finally, we measured the stability of the IRF over a two hour period starting from a cold start. This measurement characterizes the time scale of thermal equilibrium for a single Kernel Flow2 module. We analyze the total intensity, the 1st moment, and the shape of the IRF as described in the BIP protocol continuously over the two hour recording period.

#### MEDPHOT Protocol: μ_a_ and μ_s_′ measurements on optical phantoms

The MEDPHOT protocol serves as an evaluative framework for various photon migration instruments, assessing their capacity to accurately retrieve known optical properties within a physiologically relevant range using homogeneous phantoms (*23*). In this study, we employed the Kernel Flow2 modules to conduct measurements on a specific subset of the MEDPHOT kit, comprising 12 solid phantoms (BioPixS, Ireland).

These cylindrical phantoms, measuring 50 mm in height and 100 mm in diameter, consist of solid compositions containing titanium dioxide (TiO2) and absorbing toner at varying concentrations. The phantoms are identified by letters (A, B, C, and D) and numbers (1, 3, 5), where the letters denote nominal scatter values, and the numbers represent absorption values, as illustrated in Fig. 3.

Utilizing a single Flow2 module, we probed the phantoms from the top within a reflectance geometry setup. The absorption and scattering parameters were estimated by minimizing the disparity between the measured histogram and a predicted histogram, the latter being the outcome of convolving the measured impulse response function (IRF) with an analytical temporal point spread function (TPSF). The TPSF equation was derived from an analytical semi-infinite diffusion model employing Robin boundary conditions. To ensure the stability of optical property characterization, we subjected the B5 phantom to measurements for a duration of 2 hours starting from a cold start, with an independent fit of optical properties from each 1s of data.

#### nEUROPt Protocol: Depth Contrast

The nEUROPt protocol (*22*) is employed in the evaluation of devices at the system level through the utilization of optical phantoms designed to replicate brain tissue. A liquid phantom was prepared consisting of a mixture of water, India ink, and intralipid emulsion (Intralipid® 20%), titrated to have optical properties of μ_a_ = 0.01 and μ_s_′ = 1.0 mm^-1^ at 690nm. The liquid phantom tank is made of black anodized aluminum with 0.1-mm thickness mylar windows for the source and detectors, as shown in Fig. 3A. Experiments were conducted with a set of black polyvinyl chloride (PVC) cylinders with dimensions (diameter equal to height) of 3.2, 5, and 6.8mm, corresponding to volumes V_incl_/mm^3^ = 25, 100, 250. The occlusions were suspended in the titrated solution by thin metal wires (0.4mm, painted white) and were placed in the path of source-detector pairs formed within the Flow2 module. The target was moved incrementally from a depth of 8mm to 36mm away from the source-detector plane in 2mm steps. We took measurements at each depth. The original protocol calls for 100 1-s accumulation histograms, at a count rate of 1e6 photons/s. As with the BIP protocol, we matched the original protocol in two ways: with the 1s integration time (and a higher count rate), and with the target 1e6 photons per histogram (and a lower integration time).

Contrast C for a given measurand M is defined as:

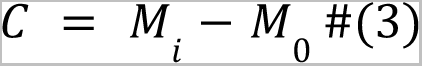

where M_0_ is the reference or baseline measurement and M_i_ us a measurement made after some i^th^ change in absorption (in our case an absorbing target) is introduced.

For photon counts, for which absolute contrast values are not as interpretable, we report a relative contrast measurement Cr defined as:

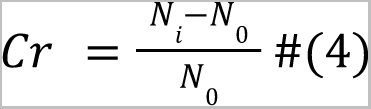

Finally, for all measurands, the contrast-to-noise ratio (CNR) is defined as:

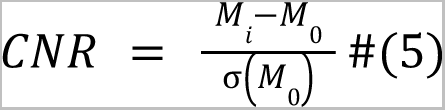

where σ(M_o_) is the standard deviation of the reference measurement across time samples. We report on these contrast metrics for 500ps time gates (after deconvolving the IRF, see next section); and for the moments of the time of flight of photons (sum, mean and variance), which are often used in TD-fNIRS analysis because they present many desirable properties (*28*).

#### Deconvolution

We can use the continuously monitored IRF and retrieve the TPSF from the DTOF. We used a FFT deconvolution approach, with a regularization factor that dampens higher frequency content (see (*57*)).

### Recordings in human participants

#### Tasks

##### Breath Hold task

This task was programmed and presented in Unity. The participant was asked to switch between interleaved periods of holding their breath and periods of paced breathing. During each paced breathing block (30 sec) a bright green circle on a black background repeatedly expanded and contracted at a fixed pace (6 sec per cycle), and the participant was instructed to use this animation to guide their inhalation and exhalation respectively (5 breathing cycles were repeated in each block). At the end of paced breathing blocks the circle changed to yellow, signaling to the participant that this would be their final exhalation and the breath hold period would occur next. During each breath hold block (20 sec) the fully contracted yellow circle remained on the screen, above which the words “Hold your breath!” were displayed. Below the yellow circle a countdown to zero indicating the time left in the block was displayed. When the countdown timer hit zero, the circle turned back to bright green and paced breathing immediately commenced. This was repeated such that the participant completed a total of 6 breath hold blocks and 6 paced breathing blocks.

##### Passive Auditory task

This task was programmed and presented in Unity. The task had a block design with two block types: story blocks (n=8) during which the participant listened to short clips from TED talks; and noise blocks (n=7) during which the participant listened to brown noise. After an initial 10s rest period, the story and noise blocks (each lasting for 20s) were presented (via earbuds) in a preset pseudo-randomized order (Fig. 6B). The participant was asked to keep their eyes open and look at a white fixation cross that was presented on a black background throughout the task.

##### Finger Tapping task

This task was programmed and presented in Unity. In this task the participant was asked to sit in a chair with their arms on the armrests such that their palms faced upwards, while audio and visual stimuli guided them through randomized periods of left and right–hand finger tapping (n = 10 blocks per side; Fig. 6E). Specifically, at the start of each block the participant was cued in two ways: 1) audibly – brown noise was played through earbuds to either the right or left ear indicating the hand that should be used during the task, and 2) visually – a white image of a hand was displayed on a black screen with either an “L” or an “R” inscribed on it, again indicating the left or right hand should be used. Both cues persisted throughout the block (17.3 sec). Within each block the participant was asked to repeatedly tap the thumb of the cued hand to a certain finger on the same hand. A red dot overlaid on a finger of the visual stimulus indicated which finger to tap. Throughout a block the red dot moved sequentially through each of the four fingers, and each shift to a new finger indicated a new trial (n = 13 trials per block; trial duration = 0.75 sec and inter-trial interval = 0.50 sec). A brief resting period (20 sec) with a white fixation cross on a black screen followed each block.

#### Data preprocessing

##### Relative changes in HbO and HbR concentrations (moments method)

The data preprocessing procedures have been extensively detailed in our previous studies (*15*). Initially, we applied a channel selection method based on histogram shape criteria (*14*). Subsequently, histograms derived from the chosen channels were utilized to calculate the moments of the DTOFs, specifically focusing on the sum, mean, and variance moments. The alterations in preprocessed DTOF moments were then translated into changes in absorption coefficients for each wavelength, employing the sensitivities of the various moments to absorption coefficient changes, as outlined in (*13*). To determine these sensitivities, a 2-layer medium with a superficial layer of 12 mm thickness was employed. Utilizing a finite element modeling (FEM) forward model from NIRFAST (*58, 59*), the Jacobians (sensitivity maps) for each moment were integrated within each layer to assess sensitivities. The changes in absorption coefficients at each wavelength were further converted into alterations in oxyhemoglobin and deoxyhemoglobin concentrations (HbO and HbR, respectively), employing the extinction coefficients for the respective wavelengths and the modified Beer–Lambert law (mBLL (*60*)). The HbO/HbR concentrations underwent additional preprocessing through a motion correction algorithm known as Temporal Derivative Distribution Repair (TDDR (*61*)). To address spiking artifacts arising from baseline shifts during TDDR, they were identified and rectified using cubic spline interpolation (*62*). Lastly, data detrending was performed using a moving average with a 100-second kernel, and short channel regression was employed to eliminate superficial physiological signals from brain activity (*63*), utilizing short within-module channels with a source-detector separation (SDS) of 8.5 mm.

##### Absolute concentrations of HbO and HbR (curve fitting method)

The DTOF results from convolving the time-resolved TPSF with the IRF. Utilizing Flow2’s online IRF measurements, we employed a curve fitting technique to extract the absolute optical properties of the tissue beneath. Generating candidate TPSFs through an analytical solution of the diffusion equation for a homogeneous semi-infinite medium, we convolved these with the known IRF and compared them with the recorded DTOF. The search for optical properties was carried out using the Levenberg-Marquardt algorithm, focusing on fitting within the range spanning from 80% of the peak on the rising edge to 0.1% of the peak on the falling edge, with a refractive index set to 1.4. These absorption coefficient estimates were then converted to HbO and HbR concentrations. A single value for HbO and HbR was obtained by computing the median value across well-coupled long, within-module channels (SDS=26.5mm) of two prefrontal modules.

##### Generalized linear model (GLM) and Epoched Analyses

A GLM approach was employed in order to elucidate patterns of significant brain activity during the different block conditions of the Auditory and Finger Tapping tasks. For each task, the activity of each channel (the hemodynamic time course, γ) was fitted with a linear model:

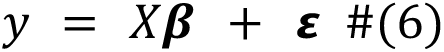

where the matrix, *X*, was composed of: i) task relevant regressors (the time course for each block condition represented as a square wave convolved with a canonical hemodynamic response function), and ii) task irrelevant regressors (namely, drift and low frequency cosine terms). A least-squares method was used to solve the multiple regression problem, producing the fitted model coefficients, **β**-weights. These weights quantify the effect that each regressor has on the hemodynamic signal. To evaluate whether the activity of a given channel is modulated across task conditions, **β**-weights associated with the block types of interest are subtracted (commonly termed as contrasts) and a t-test is employed to statistically compare this difference to zero. The contrast of interests were Story - Noise for the Auditory task and Left - Right for the Finger Tapping task. The resulting test statistics (from each channel) were plotted as a heatmap over the head to visualize patterns of brain activity and regions of interest (Fig. 6A,D).

The associated p-values from the GLM analysis were used to identify representative channels that showed significant activation to a given task condition for time course visualization and epoched analyses. These channels were chosen from the regions of interest within each task (i.e., motor and auditory areas for the Finger Tapping and Passive Auditory task respectively), and to avoid superficial signals, only channels with SDS>12 mm were considered. The time series for these channels underwent further processing; namely, detrending (with a 100 seconds kernel), and low-pass filtering (0.01-0.1Hz finite impulse response – FIR filter). For each task, the time course of HbO and HbR for one representative channel was overlaid on the task design time course (Fig. 6B,E).

Moreover, we performed standard epoching analyses by windowing the channel time courses over blocks and grouping them by block type. For both tasks the windows started 5 seconds before a block-start event and extended 30 seconds beyond (window = [-5s, 30s]; where block start = 0). A period preceding the start of each block (-5 to -1 seconds) was used to baseline the response of each epoch. The average time course (mean) and variability of response (standard error) for each block-type was computed over the condition windows (Fig. 6C,F).

##### DOT reconstruction algorithm

A finite element model (FEM) of the adult head was developed based on the unbiased non-linear averages of the MNI 152 database (*49*). The atlas was segmented to 5 tissue types of skin, skull, CSF, gray and white matter and discretized into linear tetrahedral elements using NIRFASTSlicer, giving rise to 413,403 nodes and 2,465,366 elements. Optical properties of each tissue layer at each wavelength (690 nm and 905 nm) were assigned based on published values of the adult head (*29*). The coordinates for each of the 40 modules containing the optical sources and detectors were determined and identified on the surface of the FEM and the time-resolved light propagation model was solved using the diffusion approximation to the light transport equation throughout the domain (*58*). The Jacobians (sensitivity functions that map a change in measured data due to a change in optical properties) for the time-resolved data (TPSF) for each optical parameter (μ_a_ and μ_s_′) were calculated using the adjoint theorem (*64*) at each wavelength and then interpolated to a uniform voxel grid (also known as reconstruction basis) spanning the entire model, with a resolution of 4 × 4 × 4 mm. The use of lower resolution reconstruction basis is crucial for DOT as the problem is highly under-determined: that is the number of measurements is much lower than the number of unknowns. While a high resolution FEM mesh is needed for the calculation of the time-resolved light propagation to ensure numerical accuracy, a much lower voxel resolution is needed to better improve the stability of the inverse problem.

The time-resolved Jacobian for each optical property was then mapped to each data-type (intensity, mean time of flight and variance) which was then normalized with respect to their corresponding data. A Moore–Penrose pseudoinverse with Tikhonov regularization was used to calculate an approximation of the inverse of the Jacobian to perform a single step linear recovery of the optical properties (*29*) using the same functional data as outlined earlier. Note that we downsampled the data to 1Hz before performing reconstruction. The recovered changes in the μ_a_ within each voxel were mapped to changes to oxy/deoxy hemoglobin for further processing using the same GLM model described above. Lastly, in addition to GLM analyses, we performed an epoched analysis. Here, we considered different ROIs given the task: voxels within 10 mm of the left motor area with the maximum GLM contrast for the finger tapping task and voxels within 10 mm of the left auditory region with maximum GLM contrast for the passive auditory task. The time course of these ROIs were then epoched and aggregated within each block type for further visualization (Fig. 7).

## Acknowledgments

The authors gratefully acknowledge Moriah Taylor for her administrative support for human subjects data collection, and Eli Palmer for software support with task programming.

## Author Contributions

Conceptualization: RMF, KLP.

System Design and Engineering: YC, DD, RMF, VG, IO, JS.

Software: GL, MJP, VS.

Formal analysis: JD, HD, EMK, ZMA, IO.

Investigation: VG, IO, NM.

Visualization: JD, EMK, ZMA, IO.

Writing - Original Draft: JD, HD, EMK, ZMA, IO, KLP.

Writing - Review and Editing: all authors.

Author order was determined alphabetically. Dr. Katherine Perdue agrees to be accountable for all aspects of the work, ensuring that questions related to the accuracy or integrity of any part of the work are appropriately investigated and resolved.

## References

1. S. Mattke, H. Jun, E. Chen, Y. Liu, A. Becker, C. Wallick, Expected and diagnosed rates of mild cognitive impairment and dementia in the U.S. Medicare population: observational analysis. Alz Res Therapy 15, 128 (2023).

2. World Health Organization, “World mental health report: Transforming mental health for all.” (2022).

3. M. G. Poirot, H. G. Ruhe, H.-J. M. M. Mutsaerts, I. I. Maximov, I. R. Groote, A. Bjørnerud, H. A. Marquering, L. Reneman, M. W. A. Caan, Treatment Response Prediction in Major Depressive Disorder Using Multimodal MRI and Clinical Data: Secondary Analysis of a Randomized Clinical Trial. AJP, appi.ajp.20230206 (2024).

4. M. Arns, S. Olbrich, A. T. Sack, Biomarker-driven stratified psychiatry: from stepped-care to matched-care in mental health. Nat. Mental Health 1, 917–919 (2023).

5. E. Cole, M. Gulser, K. Stimpson, B. Bentzley, J. Hawkins, X. Xiao, A. Schatzberg, K. Sudheimer, N. Williams, Stanford accelerated intelligent neuromodulation therapy for treatment-resistant depression (SAINT-TRD). Brain Stimulation 12, 402 (2019).

6. L. M. Hack, L. Tozzi, S. Zenteno, A. M. Olmsted, R. Hilton, J. Jubeir, M. S. Korgaonkar, A. F. Schatzberg, J. A. Yesavage, R. O’Hara, L. M. Williams, A Cognitive Biotype of Depression Linking Symptoms, Behavior Measures, Neural Circuits, and Differential Treatment Outcomes: A Prespecified Secondary Analysis of a Randomized Clinical Trial. JAMA Netw Open 6, e2318411 (2023).

7. W. Wu, Y. Zhang, J. Jiang, M. V. Lucas, G. A. Fonzo, C. E. Rolle, C. Cooper, C. Chin-Fatt, N. Krepel, C. A. Cornelssen, R. Wright, R. T. Toll, H. M. Trivedi, K. Monuszko, T. L. Caudle, K. Sarhadi, M. K. Jha, J. M. Trombello, T. Deckersbach, P. Adams, P. J. McGrath, M. M. Weissman, M. Fava, D. A. Pizzagalli, M. Arns, M. H. Trivedi, A. Etkin, An electroencephalographic signature predicts antidepressant response in major depression. Nat Biotechnol 38, 439–447 (2020).

8. R. Li, H. Hosseini, M. Saggar, S. C. Balters, A. L. Reiss, Current opinions on the present and future use of functional near-infrared spectroscopy in psychiatry. Neurophoton. 10 (2023).

9. H. Ayaz, W. B. Baker, G. Blaney, D. A. Boas, H. Bortfeld, K. Brady, J. Brake, S. Brigadoi, E. M. Buckley, S. A. Carp, R. J. Cooper, K. R. Cowdrick, J. P. Culver, I. Dan, H. Dehghani, A. Devor, T. Durduran, A. T. Eggebrecht, L. L. Emberson, Q. Fang, S. Fantini, M. A. Franceschini, J. B. Fischer, J. Gervain, J. Hirsch, K.-S. Hong, R. Horstmeyer, J. M. Kainerstorfer, T. S. Ko, D. J. Licht, A. Liebert, R. Luke, J. M. Lynch, J. Mesquida, R. C. Mesquita, N. Naseer, S. L. Novi, F. Orihuela-Espina, T. D. O’Sullivan, D. S. Peterka, A. Pifferi, L. Pollonini, A. Sassaroli, J. R. Sato, F. Scholkmann, L. Spinelli, V. J. Srinivasan, K. St. Lawrence, I. Tachtsidis, Y. Tong, A. Torricelli, T. Urner, H. Wabnitz, M. Wolf, U. Wolf, S. Xu, C. Yang, A. G. Yodh, M. A. Yücel, W. Zhou, Optical imaging and spectroscopy for the study of the human brain: status report. Neurophoton. 9 (2022).

10. A. Bishnoi, R. Holtzer, M. E. Hernandez, Brain Activation Changes While Walking in Adults with and without Neurological Disease: Systematic Review and Meta-Analysis of Functional Near-Infrared Spectroscopy Studies. Brain Sci 11, 291 (2021).

11. Q. Zhao, W. Zhao, C. Lu, H. Du, P. Chi, Interpersonal neural synchronization during social interactions in close relationships: A systematic review and meta-analysis of fNIRS hyperscanning studies. Neuroscience & Biobehavioral Reviews 158, 105565 (2024).

12. A. T. Eggebrecht, S. L. Ferradal, A. Robichaux-Viehoever, M. S. Hassanpour, H. Dehghani, A. Z. Snyder, T. Hershey, J. P. Culver, Mapping distributed brain function and networks with diffuse optical tomography. Nature Photon 8, 448–454 (2014).

13. A. Ortega-Martinez, D. Rogers, J. Anderson, P. Farzam, Y. Gao, B. Zimmermann, M. A. Yücel, D. A. Boas, How much do time-domain functional near-infrared spectroscopy (fNIRS) moments improve estimation of brain activity over traditional fNIRS? Neurophoton. 10 (2022).

14. H. Y. Ban, G. M. Barrett, A. Borisevich, A. Chaturvedi, J. L. Dahle, H. Dehghani, J. Dubois, R. M. Field, V. Gopalakrishnan, A. Gundran, M. Henninger, W. C. Ho, H. D. Hughes, R. Jin, J. Kates-Harbeck, T. Landy, M. Leggiero, G. Lerner, Z. M. Aghajan, M. Moon, I. Olvera, S. Park, M. J. Patel, K. L. Perdue, B. Siepser, S. Sorgenfrei, N. Sun, V. Szczepanski, M. Zhang, Z. Zhu, Kernel Flow: a high channel count scalable time-domain functional near-infrared spectroscopy system. J. Biomed. Opt. 27 (2022).

15. A. Castillo, J. Dubois, R. M. Field, F. Fishburn, A. Gundran, W. C. Ho, S. Jawhar, J. Kates-Harbeck, Z. M. Aghajan, N. Miller, K. L. Perdue, J. Phillips, W. C. Ryan, M. Shafiei, F. Scholkmann, M. Taylor, Measuring acute effects of subanesthetic ketamine on cerebrovascular hemodynamics in humans using TD-fNIRS. Sci Rep 13, 11665 (2023).

16. J. Dubois, R. M. Field, S. Jawhar, A. Jewison, E. M. Koch, Z. M. Aghajan, N. Miller, K. L. Perdue, M. Taylor, Change in brain asymmetry reflects level of acute alcohol intoxication and impacts on inhibitory control. Sci Rep 13, 10278 (2023).

17. S. R. Arridge, M. Schweiger, Image reconstruction in optical tomography. Philosophical Transactions of the Royal Society B: Biological Sciences 352, 717 (1997).

18. J. Selb, T. M. Ogden, J. Dubb, Q. Fang, D. A. Boas, Comparison of a layered slab and an atlas head model for Monte Carlo fitting of time-domain near-infrared spectroscopy data of the adult head. J. Biomed. Opt 19, 016010 (2014).

19. J. Selb, D. K. Joseph, D. A. Boas, Time-gated optical system for depth-resolved functional brain imaging. J. Biomed. Opt. 11, 044008 (2006).

20. M. A. Yücel, A. v. Lühmann, F. Scholkmann, J. Gervain, I. Dan, H. Ayaz, D. Boas, R. J. Cooper, J. Culver, C. E. Elwell, A. Eggebrecht, M. A. Franceschini, C. Grova, F. Homae, F. Lesage, H. Obrig, I. Tachtsidis, S. Tak, Y. Tong, A. Torricelli, H. Wabnitz, M. Wolf, Best practices for fNIRS publications. Neurophoton. 8 (2021).

21. H. Wabnitz, D. R. Taubert, M. Mazurenka, O. Steinkellner, A. Jelzow, R. Macdonald, D. Milej, P. Sawosz, M. Kacprzak, A. Liebert, R. Cooper, J. Hebden, A. Pifferi, A. Farina, I. Bargigia, D. Contini, M. Caffini, L. Zucchelli, L. Spinelli, R. Cubeddu, A. Torricelli, Performance assessment of time-domain optical brain imagers, part 1: basic instrumental performance protocol. J. Biomed. Opt 19, 086010 (2014).

22. H. Wabnitz, A. Jelzow, M. Mazurenka, O. Steinkellner, R. Macdonald, D. Milej, N. Zolek, M. Kacprzak, P. Sawosz, R. Maniewski, A. Liebert, S. Magazov, J. Hebden, F. Martelli, P. Di Ninni, G. Zaccanti, A. Torricelli, D. Contini, R. Re, L. Zucchelli, L. Spinelli, R. Cubeddu, A. Pifferi, Performance assessment of time-domain optical brain imagers, part 2: nEUROPt protocol. J. Biomed. Opt 19, 086012 (2014).

23. A. Pifferi, A. Torricelli, A. Bassi, P. Taroni, R. Cubeddu, H. Wabnitz, D. Grosenick, M. Möller, R. Macdonald, J. Swartling, T. Svensson, S. Andersson-Engels, Performance assessment of photon migration instruments: the MEDPHOT protocol. Applied Optics 44, 2104–2114 (2005).

24. P. Lanka, L. Yang, D. Orive-Miguel, J. D. Veesa, S. Tagliabue, A. Sudakou, S. Samaei, M. Forcione, Z. Kovacsova, A. Behera, T. Gladytz, D. Grosenick, L. Hervé, T. Durduran, K. Bejm, M. Morawiec, M. Kacprzak, P. Sawosz, A. Gerega, A. Liebert, A. Belli, I. Tachtsidis, F. Lange, G. Bale, L. Baratelli, S. Gioux, A. L. Kalyanov, M. Wolf, S. Konugolu-Venkata-Sekar, M. Zanoletti, I. Pirovano, M. Lacerenza, L. Qiu, E. Ferocino, G. Maffeis, C. Amendola, L. Colombo, L. Frabasile, P. Levoni, M. Buttafava, M. Renna, L. D. Sieno, R. Re, A. Farina, L. Spinelli, A. D. Mora, D. Contini, P. Taroni, A. Tosi, A. Torricelli, H. Dehghani, H. Wabnitz, A. Pifferi, Multi-laboratory performance assessment of diffuse optics instruments: the BitMap exercise. JBO 27, 074716 (2022).

25. R. Re, D. Contini, M. Turola, L. Spinelli, L. Zucchelli, M. Caffini, R. Cubeddu, A. Torricelli, Multi-channel medical device for time domain functional near infrared spectroscopy based on wavelength space multiplexing. *Biomed. Opt. Express*, BOE 4, 2231–2246 (2013).

26. M. Diop, K. St. Lawrence, Improving the depth sensitivity of time-resolved measurements by extracting the distribution of times-of-flight. Biomed. Opt. Express 4, 447 (2013).

27. A. Liebert, H. Wabnitz, D. Grosenick, M. Möller, R. Macdonald, H. Rinneberg, Evaluation of optical properties of highly scattering media by moments of distributions of times of flight of photons. Appl. Opt., AO 42, 5785–5792 (2003).

28. H. Wabnitz, D. Contini, L. Spinelli, A. Torricelli, A. Liebert, Depth-selective data analysis for time-domain fNIRS: moments vs time windows. Biomed. Opt. Express 11, 4224 (2020).

29. M. Doulgerakis, A. T. Eggebrecht, H. Dehghani, High-density functional diffuse optical tomography based on frequency-domain measurements improves image quality and spatial resolution. Neurophoton. 6, 035007 (2019).

30. F. Lange, I. Tachtsidis, Clinical Brain Monitoring with Time Domain NIRS: A Review and Future Perspectives. Applied Sciences 9, 1612 (2019).

31. J. Selb, J. J. Stott, M. A. Franceschini, A. G. Sorensen, D. A. Boas, Improved sensitivity to cerebral hemodynamics during brain activation with a time-gated optical system: analytical model and experimental validation. J. Biomed. Opt. 10, 011013 (2005).

32. M. M. Wu, K. Perdue, S.-T. Chan, K. A. Stephens, B. Deng, M. A. Franceschini, S. A. Carp, Complete head cerebral sensitivity mapping for diffuse correlation spectroscopy using subject-specific magnetic resonance imaging models. Biomed. Opt. Express 13, 1131 (2022).

33. U. E. Emir, C. Ozturk, A. Akin, Multimodal investigation of fMRI and fNIRS derived breath hold BOLD signals with an expanded balloon model. Physiol. Meas. 29, 49–63 (2008).

34. J. Dubois, R. M. Field, S. Jawhar, E. M. Koch, Z. M. Aghajan, N. Miller, K. L. Perdue, M. Taylor, “Reliability of brain metrics derived from a Time-Domain Functional Near-Infrared Spectroscopy System” (preprint, BioRxiv, 2024); 10.1101/2024.03.12.584660.

35. B. J. MacIntosh, L. M. Klassen, R. S. Menon, Transient hemodynamics during a breath hold challenge in a two part functional imaging study with simultaneous near-infrared spectroscopy in adult humans. NeuroImage 20, 1246–1252 (2003).

36. I. Tonhajzerova, M. Mestanik, A. Mestanikova, A. Jurko, Respiratory sinus arrhythmia as a non-invasive index of ′brain-heart′ interaction in stress. Indian J Med Res 144, 815 (2016).

37. P. Belin, R. J. Zatorre, P. Lafaille, P. Ahad, B. Pike, Voice-selective areas in human auditory cortex. Nature 403, 309–312 (2000).

38. R. Luke, E. Larson, M. J. Shader, H. Innes-Brown, L. Van Yper, A. K. C. Lee, P. F. Sowman, D. McAlpine, Analysis methods for measuring passive auditory fNIRS responses generated by a block-design paradigm. Neurophoton. 8 (2021).

39. J. Brinkman, H. G. J. M. Kuypers, Cerebral control of contralateral and ipsilateral arm, hand and finger movements in the split-brain rhesus monkey. Brain 96, 653–674 (1973).

40. H. Obrig, J. Steinbrink, Non-invasive optical imaging of stroke. *Philosophical Transactions of the Royal Society A: Mathematical*, Physical and Engineering Sciences, doi: 10.1098/rsta.2011.0252 (2011).

41. R. Thomas, S. S. Shin, R. Balu, Applications of near-infrared spectroscopy in neurocritical care. Neurophotonics 10 (2023).

42. A. Abdalmalak, D. Milej, M. Diop, M. Shokouhi, L. Naci, A. M. Owen, K. St. Lawrence, Can time-resolved NIRS provide the sensitivity to detect brain activity during motor imagery consistently? Biomed. Opt. Express 8, 2162 (2017).

43. R. Yang, J. F. Dunn, Reduced cortical microvascular oxygenation in multiple sclerosis: a blinded, case-controlled study using a novel quantitative near-infrared spectroscopy method. Sci Rep 5, 16477 (2015).

44. Y. Hoshi, T. Shinba, C. Sato, N. Doi, Resting hypofrontality in schizophrenia: A study using near-infrared time-resolved spectroscopy. Schizophrenia Research 84, 411–420 (2006).

45. S. Viola, P. Viola, M. P. Buongarzone, L. Fiorelli, P. Litterio, Tissue oxygen saturation and pulsatility index as markers for amnestic mild cognitive impairment: NIRS and TCD study. Clinical Neurophysiology 124, 851–856 (2013).

46. S. T. Witt, A. R. Laird, M. E. Meyerand, Functional neuroimaging correlates of finger-tapping task variations: An ALE meta-analysis. NeuroImage 42, 343–356 (2008).

47. E. E. Vidal-Rosas, A. von Lühmann, P. Pinti, R. J. Cooper, Wearable, high-density fNIRS and diffuse optical tomography technologies: a perspective. NPh 10, 023513 (2023).

48. I. Mazzonetto, M. Castellaro, R. J. Cooper, S. Brigadoi, Smartphone-based photogrammetry provides improved localization and registration of scalp-mounted neuroimaging sensors. Sci Rep 12, 10862 (2022).

49. V. Fonov, A. Evans, R. McKinstry, C. Almli, D. Collins, Unbiased nonlinear average age-appropriate brain templates from birth to adulthood. NeuroImage 47, S102 (2009).

50. F. Gao, H. Zhao, Y. Yamada, Improvement of image quality in diffuse optical tomography by use of full time-resolved data. *Appl. Opt.*, AO 41, 778–791 (2002).

51. M. D. Wheelock, J. P. Culver, A. T. Eggebrecht, High-density diffuse optical tomography for imaging human brain function. Review of Scientific Instruments 90, 051101 (2019).

52. A. Bonilauri, F. Sangiuliano Intra, L. Pugnetti, G. Baselli, F. Baglio, A Systematic Review of Cerebral Functional Near-Infrared Spectroscopy in Chronic Neurological Diseases—Actual Applications and Future Perspectives. Diagnostics 10, 581 (2020).

53. H. Hampel, P. Gao, J. Cummings, N. Toschi, P. M. Thompson, Y. Hu, M. Cho, A. Vergallo, The foundation and architecture of precision medicine in neurology and psychiatry. Trends in Neurosciences 46, 176–198 (2023).

54. S. Nebe, M. Reutter, D. H. Baker, J. Bölte, G. Domes, M. Gamer, A. Gärtner, C. Gießing, C. Gurr, K. Hilger, P. Jawinski, L. Kulke, A. Lischke, S. Markett, M. Meier, C. J. Merz, T. Popov, L. M. Puhlmann, D. S. Quintana, T. Schäfer, A.-L. Schubert, M. F. Sperl, A. Vehlen, T. B. Lonsdorf, G. B. Feld, Enhancing precision in human neuroscience. eLife 12, e85980 (2023).

55. S. L. Rossi, P. Subramanian, D. E. Bovenkamp, The future is precision medicine-guided diagnoses, preventions and treatments for neurodegenerative diseases. Front. Aging Neurosci. 15, 1128619 (2023).

56. M. Lacerenza, M. Buttafava, M. Renna, A. D. Mora, L. Spinelli, F. Zappa, A. Pifferi, A. Torricelli, A. Tosi, D. Contini, Wearable and wireless time-domain near-infrared spectroscopy system for brain and muscle hemodynamic monitoring. Biomed. Opt. Express 11, 5934 (2020).

57. M. F. Wahab, T. C. O’Haver, Peak deconvolution with significant noise suppression and stability using a facile numerical approach in Fourier space. Chemometrics and Intelligent Laboratory Systems 235, 104759 (2023).

58. H. Dehghani, M. E. Eames, P. K. Yalavarthy, S. C. Davis, S. Srinivasan, C. M. Carpenter, B. W. Pogue, K. D. Paulsen, Near infrared optical tomography using NIRFAST: Algorithm for numerical model and image reconstruction. Commun Numer Methods Eng 25, 711–732 (2008).

59. M. Doulgerakis-Kontoudis, A. T. Eggebrecht, S. Wojtkiewicz, J. P. Culver, H. Dehghani, Toward real-time diffuse optical tomography: accelerating light propagation modeling employing parallel computing on GPU and CPU. JBO 22, 125001 (2017).

60. T. J. Huppert, S. G. Diamond, M. A. Franceschini, D. A. Boas, HomER: a review of time-series analysis methods for near-infrared spectroscopy of the brain. Appl. Opt. 48, D280 (2009).

61. F. A. Fishburn, R. S. Ludlum, C. J. Vaidya, A. V. Medvedev, Temporal Derivative Distribution Repair (TDDR): A motion correction method for fNIRS. NeuroImage 184, 171–179 (2019).

62. F. Scholkmann, S. Spichtig, T. Muehlemann, M. Wolf, How to detect and reduce movement artifacts in near-infrared imaging using moving standard deviation and spline interpolation. Physiol. Meas. 31, 649–662 (2010).

63. L. Gagnon, K. Perdue, D. N. Greve, D. Goldenholz, G. Kaskhedikar, D. A. Boas, Improved recovery of the hemodynamic response in diffuse optical imaging using short optode separations and state-space modeling. NeuroImage 56, 1362–1371 (2011).

64. S. R. Arridge, M. Schweiger, Photon-measurement density functions Part 2: Finite-element-method calculations. Appl. Opt. 34, 8026 (1995).

