## supplementary material for "Time-Domain Diffuse Optical Tomography for Precision Neuroscience"

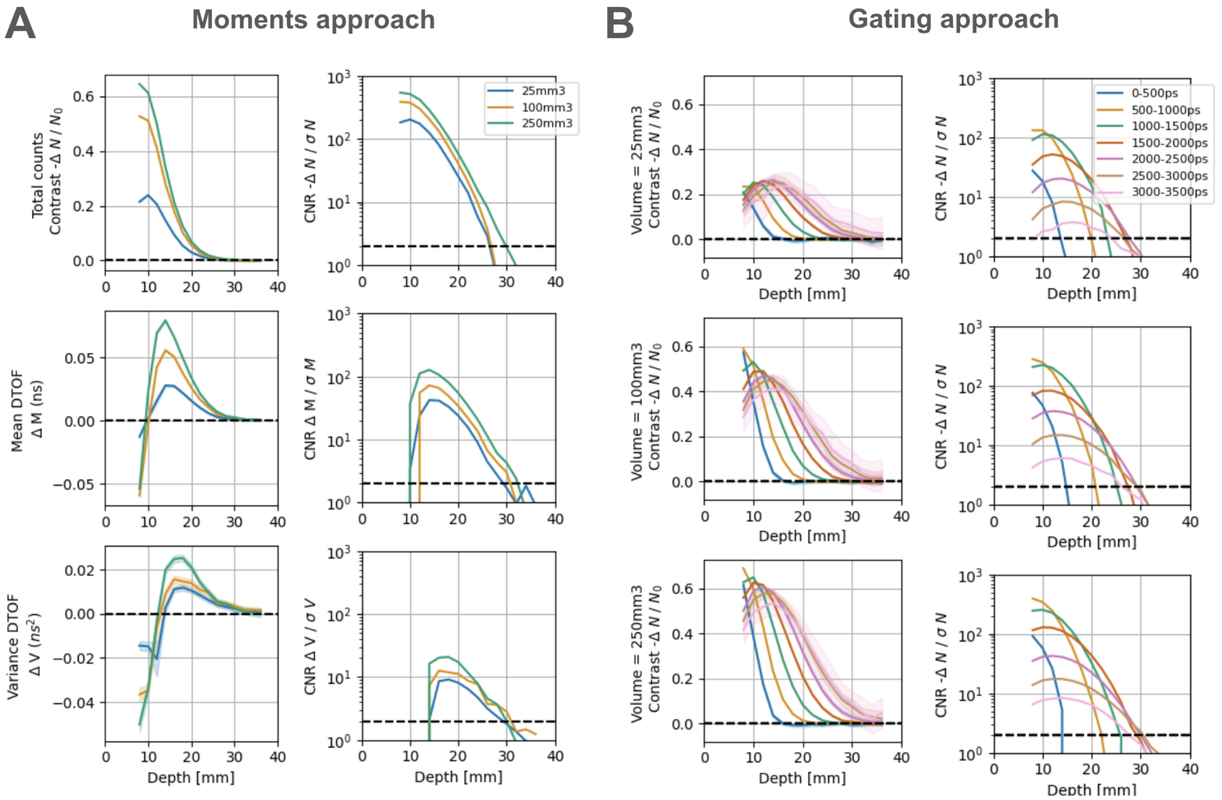


**Fig. S1. nEUROPt protocol results for 690 nm wavelength. (A)** Left) Contrast as a function of depth for three different moments of DTOFs (sum, mean and variance are the three rows respectively). Each color represents the volume of a different PVC cylinder used in the experiment. Right) Same as (left) but the contrast-to-noise ratio (CNR) at different depths. **(B)** Contrast (left) and CNR (right) are shown as a function of depth for the three different PVC cylinders used (each row). Each color corresponds to different time gates used for analysis. Both A and B are results with the 690 nm wavelength and a representative channel. For the results with 905 nm wavelength see Fig. 4.

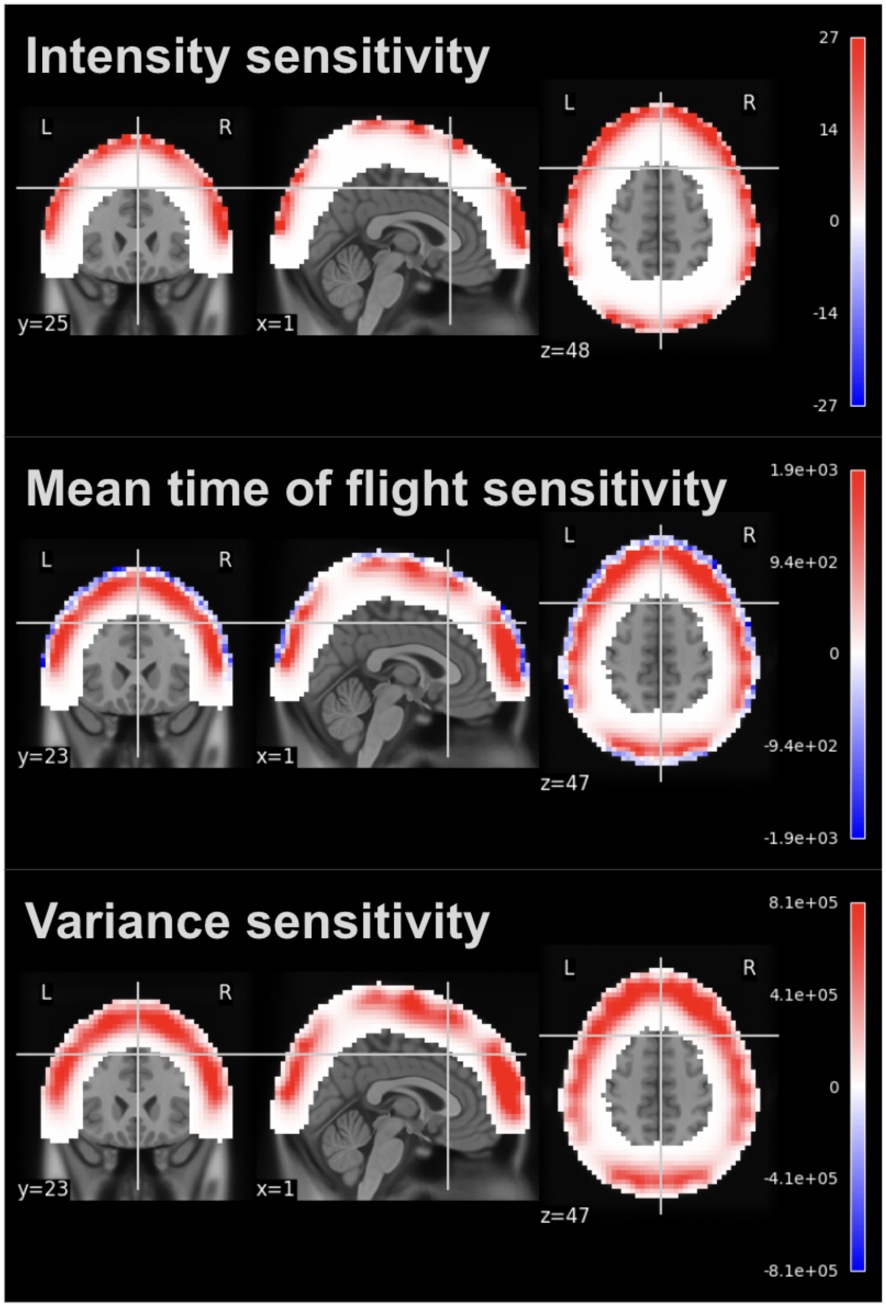


**Fig. S2. Sensitivity profiles for intensity, mean time of flight, and variance moments.** The computed Jacobians are shown for the first three moments (each row) in the MNI space using the coronal (left), sagittal (middle), and axial (right) views. Note that higher moments exhibit sensitivity to deeper layers as indicated by the thicker red regions. The areas with lower sensitivity in the superficial layers are due to the gaps between the headset plates and lower number of channels in those regions (e.g., the area between the prefrontal and frontoparietal plate).
